# Dominant toxicity of ALS–FTD-associated *CHCHD10^S59L^* is mediated by TDP-43 and PINK1

**DOI:** 10.1101/753558

**Authors:** Minwoo Baek, Yun-Jeong Choe, Gerald W. Dorn, J. Paul Taylor, Nam Chul Kim

**Affiliations:** Department of Pharmacy Practice and Pharmaceutical Sciences, College of Pharmacy, University of Minnesota, Duluth, MN 55812, USA; Center for Pharmacogenomics, Washington University School of Medicine, St. Louis, MO 63130, USA; Howard Hughes Medical Institute and Department of Cell and Molecular Biology, St. Jude Children’s Research Hospital, Memphis, TN 38105, USA

## Abstract

Mutations in coiled-coil-helix-coiled-coil-helix domain containing 10 (*CHCHD10*) are a genetic cause of amyotrophic lateral sclerosis and/or frontotemporal dementia (ALS-FTD). To elucidate how mutations in *CHCHD10* induce disease, we generated a *Drosophila melanogaster* model of *CHCHD10*-mediated ALS-FTD. Expression of *CHCHD10^S59L^* in *Drosophila* caused gain-of-function toxicity in eyes, motor neurons, and muscles, in addition to mitochondrial defects in flies and HeLa cells. TDP-43 and PINK1 formed two axes, driving the mutant-dependent phenotypes. *CHCHD10^S59L^* expression increased TDP-43 insolubility and mitochondrial translocation. Blocking mitochondrial translocation with a peptide inhibitor reduced *CHCHD10^S59L^*-mediated toxicity. *PINK1* knockdown rescued *CHCHD10^S59L^*-mediated phenotypes in *Drosophila* and HeLa cells. The two PINK1 substrates mitofusin and mitofilin were genetic modifiers of this phenotype. Mitofusin agonists reversed the *CHCHD10^S59L^*-induced phenotypes in *Drosophila* and HeLa cells and increased ATP production in *Drosophila* expressing *C9orf72* with expanded GGGGCC repeats. Two peptides inhibitors of PINK1 mitigated the mitochondrial defects introduced by *CHCHD10^S59L^* expression. These findings indicate that TDP-43 mitochondrial translocation and chronic activation of PINK1-mediated pathways by CHCHD10^S59L^ generate dominant toxicity. Therefore, inhibiting PINK1 activity may provide a therapeutic strategy for *CHCHD10*-associated disease.

**One Sentence Summary:** Inhibition of TDP-43 mitochondrial translocation or PINK1 kinase activity mitigates *CHCHD10^S59L^*-mediated mitochondrial toxicity.

## Introduction

In 2014, Bannwarth *et al*. identified an S59L substitution in coiled-coil-helix-coiled-coil-helix domain containing 10 (CHCHD10) as the causes of a familial disease characterized by motor function defects, declined cognitive function, and myopathy (*1*). Subsequent analyses revealed that *CHCHD10^S59L^* is a genetic cause of amyotrophic lateral sclerosis with frontotemporal dementia (ALS-FTD) (*1*). *CHCHD10* encodes a functionally unknown, small protein comprising a putative *N*-terminal mitochondrial–targeting sequence and a *C*-terminal CHCHD domain (*1*).

Many additional *CHCHD10* variants are now known to cause ALS, FTD, and other related degenerative diseases (*2, 3*). However, their pathogenicity and penetrance are debatable. Although *CHCHD10^R15L^* was identified in familial and sporadic cases of ALS, the existence of unaffected carriers in familial cases suggests incomplete penetrance (*4–7*). The *CHCHD10^P34S^* variant occurs in sporadic ALS (*8, 9*), ALS-FTD (*10*), Parkinson disease (*6*), and Alzheimer disease (*6*), and its overexpression causes mitochondrial defects (*11*). However, its pathogenicity is not well supported by genetic evidence (*12, 13*). *CHCHD10^G58R^* and *CHCHD10^G66V^* were identified in mitochondrial myopathy and late-onset spinal muscular atrophy, jokela type (SMAJ) or Charcot-Marie-Tooth disease type 2 (CMT2), respectively (*5, 14–16*). Other ALS–FTD-associated genes, such as valosin-containing protein (*VCP*) and Matrin 3 (*MATR3*), also exhibit clinical pleiotropy, including myopathy (*17, 18*).

*CHCHD10* mutations identified in familial diseases are dominantly inherited (*1, 4, 5, 14*). However, experimental evidence does not support that all disease-causing variants have the same mode of action. *CHCHD10* expression in patient tissues is unaffected, and *CHCHD10^S59L^* overexpression causes mitochondrial defects similar to those in affected patients (*1*). This suggests that *CHCHD10^S59L^* is a dominant gain-of-function mutation. However, *CHCHD10^S59L^* does not retain its wild-type (WT)-like activity, indicative of a dominant-negative mechanism (*19*). Furthermore, patient fibroblasts with either *CHCHD10^R15L^* or *CHCHD10^G66V^* exhibit reduced expression and protein instability, supporting a haploinsufficiency mechanism (*20, 21*).

Although some molecular mechanisms for *CHCHD10*-mediated toxicity are known, it is unclear how these mechanisms drive the disease phenotype and whether they can be controlled therapeutically. CHCHD10 interacts with components of the mitochondrial contact site and cristae organizing system (MICOS), and MICOS is decreased in patients with *CHCHD10* mutations (*11*). However, CHCHD10 is not well localized with MICOS, and CHCHD10– CHCHD2 hetero-complex formation decreases in patient fibroblasts carrying *CHCHD10^R15L^* (*21*). Although CHCHD10 and TDP-43 physically interact (*19*), phosphorylated TDP-43 levels are not associated with the phenotypic severity of *CHCHD10^S59L^* or *CHCHD10^G66V^* (*22*). Indeed, severity is more closely associated with MICOS formation (*22*).

Identification of *CHCHD10* mutations and mitochondrial pathogenic pathways in ALS-FTD suggest that mitochondrial defects are a primary cause of ALS, FTD, or other related diseases (*23–28*). To address the role of mitochondrial defects in ALS-FTD pathogenesis, we used *Drosophila* and mammalian cell culture models of mutant *CHCHD10*-mediated toxicity. *CHCHD10^S59L^* expression imparted a toxic gain of function that was mediated through two distinct axes: TDP-43 and PINK1. Pharmacologic treatment with peptide inhibitors of TDP-43 mitochondrial translocation or PINK1 kinase activity mitigated degenerative phenotypes in HeLa cells. Small-molecule agonists of mitofusin (MFN), a downstream substrate of PINK1, rescued mutant *CHCHD10*-induced mitochondrial defects in *Drosophila* and HeLa cells and enhanced mitochondrial ATP production in a *Drosophila* ALS-FTD model expressing *C9orf72* with expanded GGGGCC repeats.

## Results

### CHCHD2 and CHCHD10 share a common Drosophila melanogaster ortholog

To elucidate whether an ortholog of *CHCHD10* exists in *Drosophila*, we used the *Drosophila* RNAi Screening Center’s Integrative Ortholog Prediction Tool (*29*) and found that *Drosophila* and humans have three and two homologous genes for *CHCHD10*, respectively. Further phylogenetic analysis with a neighbor-joining tree of these genes revealed that *Dmel\CG5010* shares the highest homology with human *CHCHD2* and *CHCHD10* (fig. S1A). Two additional *Drosophila* homologues, *Dmel\CG31007* and *Dmel\CG31008*, appeared independently after their speciation. The substituted amino acids in humans are conserved in *Dmel\CG5010* (Fig. 1A). *Dmel\CG5010* is strongly expressed in all *Drosophila* tissues, whereas *Dmel\CG31007* and *Dmel\CG31008* are expressed only in the testis and weakly in the imaginal discs. A comparison between *H. sapiens* and *M. musculus* suggested that *CHCHD10* and *CHCHD2* were duplicated before their speciation and are involved in common processes, but in a distinct manner (*30*). On the basis of its phylogenetics and expression profile, we deemed *Dmel\CG5010* as a common *Drosophila* ortholog for both *CHCHD10* and *CHCHD2*. Therefore, we henceforth refer to *Dmel\CG5010* as *C2C10H* (i.e., *CHCHD2* and *CHCHD10* homolog).

**Fig 1.**
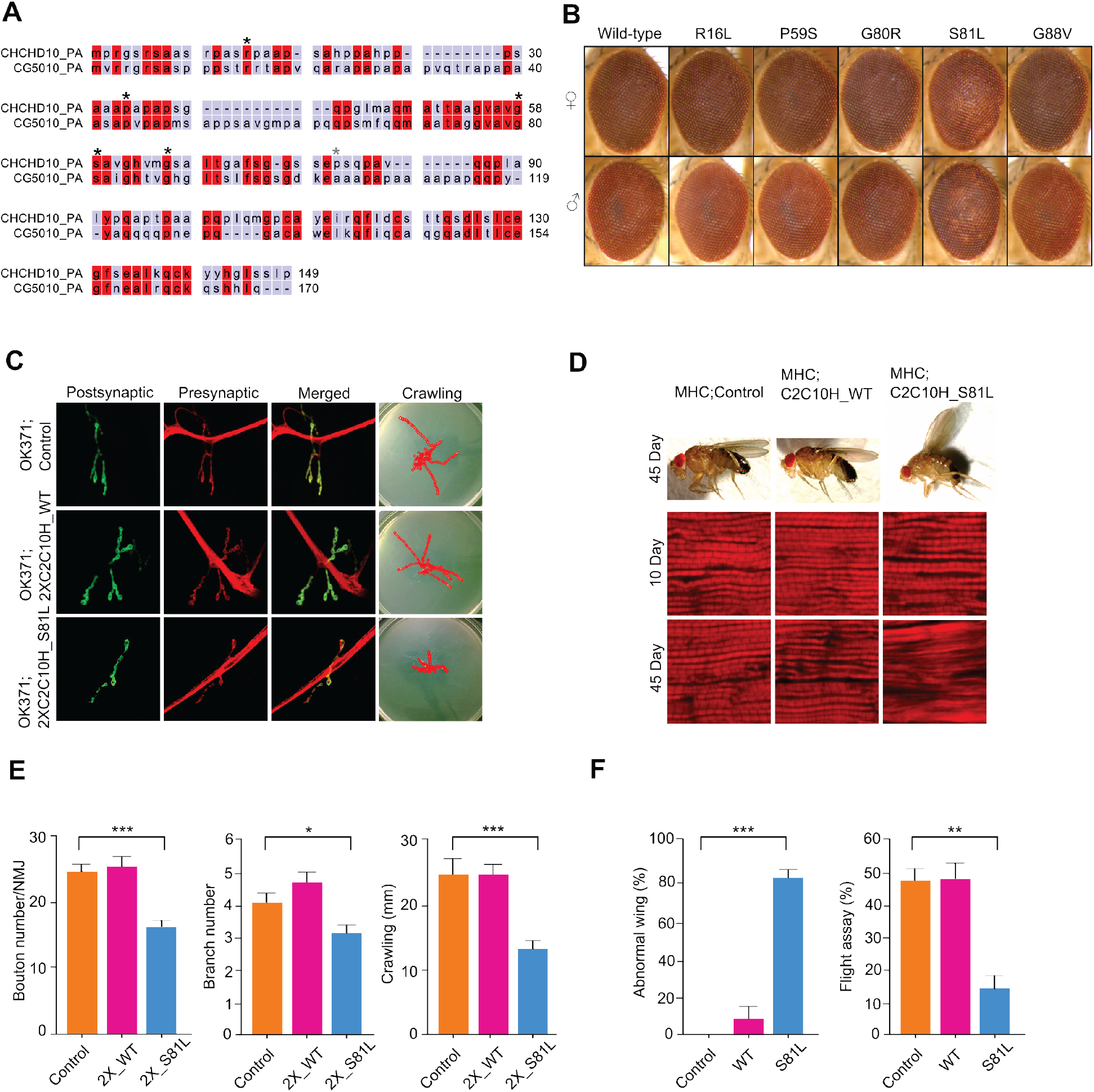
*C2C10H^S81L^* is toxic in *Drosophila* eyes, neurons, and muscles. (**A**) Protein sequence alignment of human CHCHD10 and *Drosophila* C2C10H (CG5010). Disease-causing sites (asterisk) are conserved between human CHCHD10 and *Drosophila* C2C10H. (**B**) *C2C10H^S81L^* causes age-dependent rough eye phenotypes. (**C**) Representative images of neuromuscular junctions and crawling traces from the genotypes indicated. (**D**) Adult thoraxes dissected to expose longitudinal indirect flight muscles and stained with phalloidin–Alexa Fluor 594. Flies expressing *C2C10H^S81L^* in muscles under control of *MHC*-GAL4 exhibit disrupted sarcomere structures. (**E**) Expression of *C2C10H^S81L^* in motor neurons results in small synapses, with reduced bouton and branch numbers and defective locomotor activity assessed by the crawling behavior of third-instar larvae. Data are mean ± SE (one-way ANOVA, *\*P < 0.05, ***P < 0.001*). (**F**) Expression of *C2C10H^S81L^* in muscle tissues causes abnormal wing postures and locomotor defects assessed by flight ability. Data are mean ± SE (one-way ANOVA, \*\**P* < 0.01, \*\*\**P* < 0.001; *n* = 4).

### CHCHD10^S59L^ causes dominant toxicity in fly eyes, but other mutants do not

To develop *Drosophila* models for mutant *CHCHD10*-induced human disease, we generated a series of transgenic fly lines carrying WT and mutated *CHCHD10* cDNA under the control of the upstream activating sequence (*UAS*) that expresses a downstream transgene combined with a tissue-specific GAL4 driver. We generated both *Drosophila* and codon-optimized human versions of transgenic animals carrying *CHCHD10^WT^* and the R15L, P34S, G58R, S59L, and G66V substitutions individually (i.e., R16L, P59S, G80R, S81L, and G88V in *Drosophila*, respectively), with or without a *C-*terminal FLAG tag by ΦC31 integrase-mediated site-specific integration into the *attp2* landing site on the third chromosome. Because the pathogenicity of CHCHD10^P34S^ is not well supported and controversial, we included the P34S substitution to examine its pathogenicity.

When *C2C10H^WT^* and the variants were expressed in *Drosophila* eyes by the *glass* multimer reporter (*GMR*)-GAL4 driver, they did not cause any abnormal phenotypes at eclosion. However, expression of *C2C10H^S81L^* (i.e., the fly homolog of *CHCHD10^S59L^*) caused mild but mutation-dependent depigmentation as the flies aged, whereas the other mutants (both *Drosophila* and human orthologs) did not cause any notable phenotypes by 40 days, regardless of FLAG tagging (Fig. 1B and fig. S1B and C). The expression ratio of exogenous *C2C10H* to endogenous *C2C10H* was unremarkable compared to that of our previously reported overexpression model of the *Drosophila* ortholog of *VCP* (*TER94*), which exhibits a robust rough eye phenotype at eclosion (fig. S1D and E). When we generated fly lines carrying two copies of the *C2C10H* variants at the *attp2* locus with *GMR*-GAL4, the rough eye phenotype of *C2C10H^S81L^* was enhanced and obvious at eclosion due to high levels of expression by transvection (fig. S1F) (*31*). However, expression of *C2C10H^WT^* and the variants did not induce any eye defects. The difference among the transgenes was evidenced by lethality, which was caused by leaky expression of *GMR*-GAL4 in other tissues (*32*). Homozygous flies with both *GMR*-GAL4 and either *C2C10H^WT^* or *C2C10H^P59S^* were viable and fertile. In contrast, flies expressing *C2C10H^R16L^* with *GMR*-GAL4 had reduced levels of homozygote formation. Very few homozygous flies carrying *C2C10H^G80R^* or *C2C10H^G88V^* were detected across several generations. Homozygous *C2C10H^S81L^* flies were relatively healthy and fertile but exhibited severe eye defects. This observation implies that the toxicity of each variant manifests differently in different tissues, recapitulating variant expression in clinically similar but different diseases.

*C2C10H^S81L^ recapitulates morphologic and functional defects in motor neurons and muscles* Patients with *CHCHD10^S59L^*-mediated disease experience motor neuron defects and myopathy. We next examined whether *C2C10H^S81L^* expression causes degenerative phenotypes in *Drosophila* motor neurons and muscles. Expressing *C2C10H^WT^* or *C2C10H^S81L^* in motor neurons with the motor neuron–specific driver *OK371*-GAL4 did not cause any abnormalities in larvae or adult flies, including viability and fertility. However, homozygous animals for *OK371*-GAL4 and *C2C10H^S81L^* exhibited robust degenerative phenotypes, whereas homozygous *C2C10H^WT^* flies with *OK371*-GAL4 did not show any abnormalities. Third-instar *C2C10H^S81L^* homozygous larvae showed striking locomotor dysfunction (Fig. 1C and E), with marked morphologic defects in their neuromuscular junctions (NMJs), including decreased synaptic bouton and branch numbers. Such NMJ defects were absent in driver-only or *C2C10H^WT^*-expressing flies (Fig 1. C and E). Expression of *C2C10H^S81L^* but not *C2CH10^WT^* in indirect flight muscles with the muscle-specific driver *MHC*-GAL4 caused mutant-dependent muscle degeneration, which was characterized by abnormal wing posture and loss of sarcomere architecture in aged flies (Fig. 1D and F). Additionally, *C2C10H^S81L^* expression in muscles resulted in functional locomotor defects, as evident in flight assays (Fig. 1F). Therefore, the *Drosophila* model recapitulates the dominant toxicity of *CHCHD10^S59L^ in vivo*.

### CHCHD10 mutants induce mitochondrial defects

CHCHD10 is primarily localized at cristae junctions in the mitochondrial intermembrane space (*1, 33*). To determine whether the *CHCHD10* variants induce mitochondrial defects, we transiently expressed *CHCHD10^WT^* or mutants (i.e., R15L, P34S, G58R, S59L, and G66V) tagged with *C*-terminal FLAG in HeLa cells. All variants were expressed at similar steady-state levels (fig. S2), and CHCHD10 was localized in mitochondria but not in other cellular sites (Fig. 2A). Consistent with previous reports, expression of *CHCHD10^S59L^* caused the most notable mitochondrial fragmentation and functional respiratory defects. In contrast, expression of *CHCHD10^WT^* induced mitochondrial elongation and improved respiratory function over that of empty vector–transfected cells (Fig. 2A–C). Expression of *CHCHD10^P34S^* did not cause any obvious morphologic or functional defects in HeLa cells, and expression of *CHCHD10^R15L^* resulted in mild, but statistically significant (*P* = 0.0217) morphologic defects without functional respiratory defects (Fig. 2A–C), which is discordant with our findings in *Drosophila*. CHCHD10^S59L^ displayed a punctate staining pattern in mitochondria, whereas CHCHD10^WT^ staining was evenly distributed (Fig. 2A and B). Expression of *CHCHD10^G58R^* or *CHCHD10^G66V^* also resulted in mitochondrial fragmentation and a punctate staining pattern comparable to that of CHCHD10^S59L^ (Fig. 2A and B). It is unclear whether this punctate pattern is due to CHCHD10 aggregation or mitochondrial fragmentation from conventional confocal microscopy because the staining patterns of the mitochondrial marker and CHCHD10 mostly overlapped (Fig. 2A and B).

**Fig. 2.**
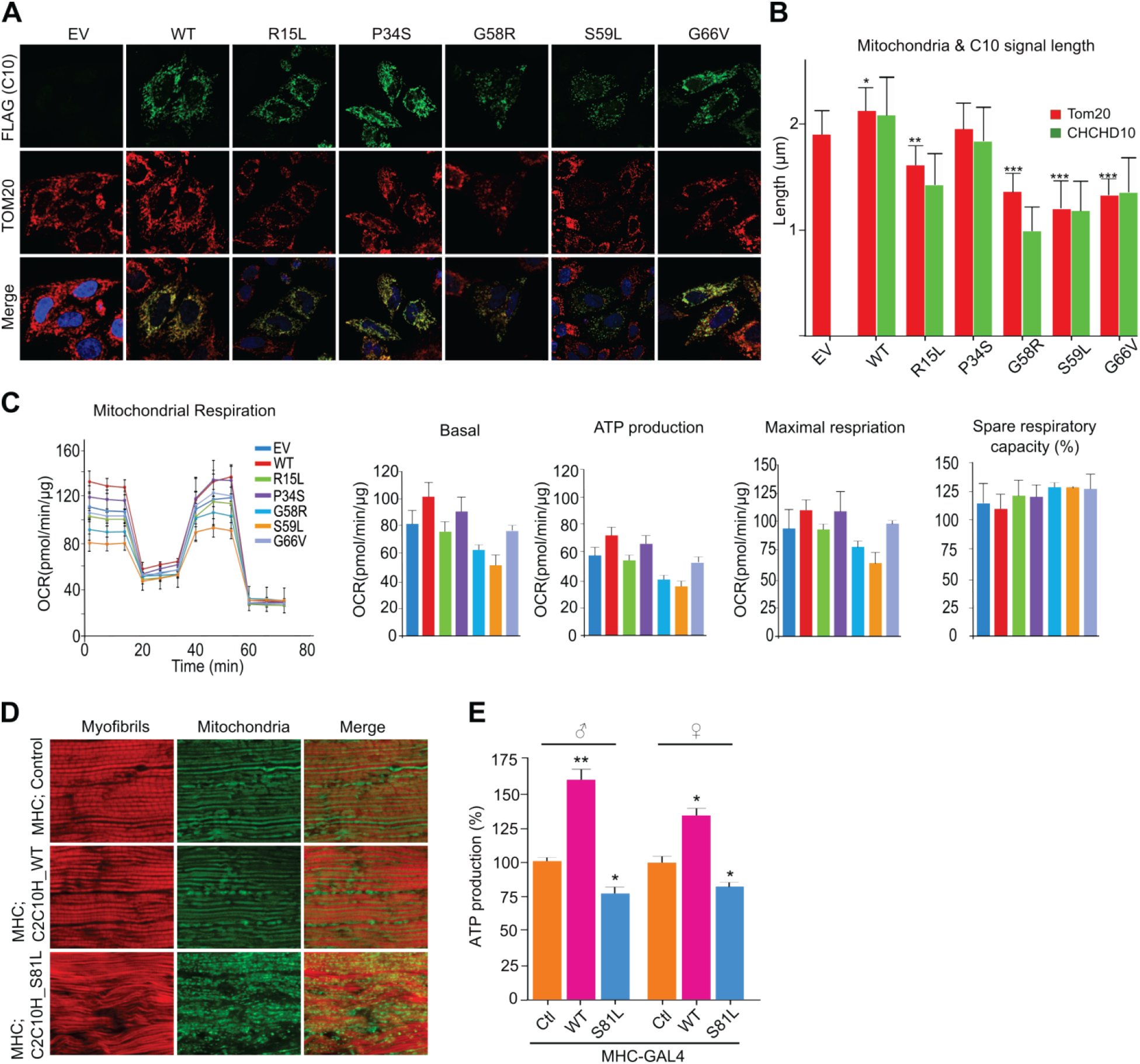
*CHCHD10* mutants induce mitochondrial defects. HeLa cells were transfected with FLAG-tagged *CHCHD10 ^WT^*, mutants (R15L, P34S, G58R, S59L, G66V), or empty vector (EV). (**A**) Representative images of HeLa cells immunostained with antibodies against FLAG (green, transfected *CHCHD10*) and TOM20 (red, mitochondria) 24 hours after transfection. (**B**) Quantification of FLAG-CHCHD10 (green) and TOM20 (red) signal strength. Data shown are mean ± SD (one-way ANOVA followed by Tukey posthoc comparison with EV, \**P* < 0.05, \*\**P* < 0.01, \*\*\**P* < 0.001; *n* = 3 biological replicates, with ≥ 20 cells measured in triplicate for each sample). (**C**) Oxygen consumption rates for each transfectant were measured. After measuring basal respiration rates, oligomycin, FCCP, and antimycin A/rotenone were serially injected to measure ATP production, maximal respiration, and non-mitochondrial respiration, respectively. Spare respiratory capacity was then calculated. Data shown are mean ± SD (*n* = 3 technical replicates). (**D**) Mitochondrial morphology of 45-day-old *C2C10H^WT^*-and *C2C10H^S81L^*-expressing flies. Indirect flight muscles were stained with streptavidin–Alexa Fluor 488 (mitochondria, green) and phalloidin–Alexa Fluor 594 (actin filaments, red). (**E**) ATP levels in thoraxes containing muscle tissues of the indicated genotypes (aged 10 days) were measured. ATP levels were normalized to total protein concentrations. Data are mean ± SE (one-way ANOVA, \**P* < 0.05, \*\**P* < 0.01; *n* = 3 replicates).

To examine mitochondrial morphology and function in response to *C2C10H^S81L^* expression in *Drosophila*, we expressed two copies of *C2C10H^S81L^* in muscle tissues with the *MHC*-GAL4 driver. Expression of *C2C10H^S81L^* in indirect flight muscles resulted in muscular degeneration and fragmented mitochondria, in contrast with that in the indirect flight muscles of *MHC*-GAL4 only or *C2C10H^WT^*-expressing flies (Fig. 2D). We measured ATP levels as an indicator of mitochondrial dysfunction in the thoraxes of the flies, which contain primarily muscle tissue. Significantly reduced (*P* = 0.0235) ATP levels occurred in the muscle tissues of flies expressing *C2C10H^S81L^*, as compared with those of *MHC*-GAL4/+ control flies (Fig. 2E). Consistent with the effect of *CHCHD10^WT^* overexpression in HeLa cells, *C2C10H^WT^* expression also increased ATP levels in *Drosophila* muscle tissues (Fig. 2E). Therefore, only *CHCHD10^S59L^* induced consistent mitochondrial fragmentation and functional respiratory defects in both *Drosophila* and mammalian cells.

### CHCHD10^S59L^-induced phenotypes are rescued by WT and mutant co-expression

The S59L substitution in CHCHD10 is dominantly inherited (*1*). The dominant toxicity imparted by *CHCHD10^S59L^* overexpression suggests two possible modes of action: dominant negative or dominant gain of function. We generated a fly model carrying two copies of *C2C10H^S81L^* in the second and the third chromosomes. When the two *C2C10H^S81L^* copies were expressed via *GMR*-GAL4, it induced a relatively mild rough eye phenotypes at eclosion (Fig. 3A), as compared with that of third chromosome homozygotes (fig. S1D). *C2C10H^WT^* co-expression suppressed the *C2C10H^S81L^*-induced rough eye phenotype (Fig. 3A), in addition to the morphologic and functional mitochondrial defects in both *Drosophila* (Fig. 3B and C) and HeLa cells (Fig. 3D and E and fig. S3A). This suggests that *CHCHD10^S59L^* is a dominant-negative mutant. To exclude the possibility of unknown positional effects such as transvection, we generated another transgenic fly line with *C2C10H^WT^* or *C2C10H^S81L^* in an unrelated landing site, *VK27*. *C2C10H^WT^_VK27_* also rescued the homozygous *C2C10H^S81L^* phenotypes, whereas *C2C10H^S81L^_VK27_* enhanced the rough eye phenotype (fig. S3B). Surprisingly, all co-expressed variants reduced the rough eye phenotypes with subtle differences (Fig. 3A and fig. S3C). Co-expression of *C2C10H^WT^* and each of the variants also mitigated *C2C10H^S81L^*-induced mitochondrial fragmentation and ATP production defects in indirect flight muscles (Fig. 3B and C). An additional copy of *C2C10H^S81L^* exacerbated the defects in both eyes and muscles (Fig. 3A–C).

**Fig. 3.**
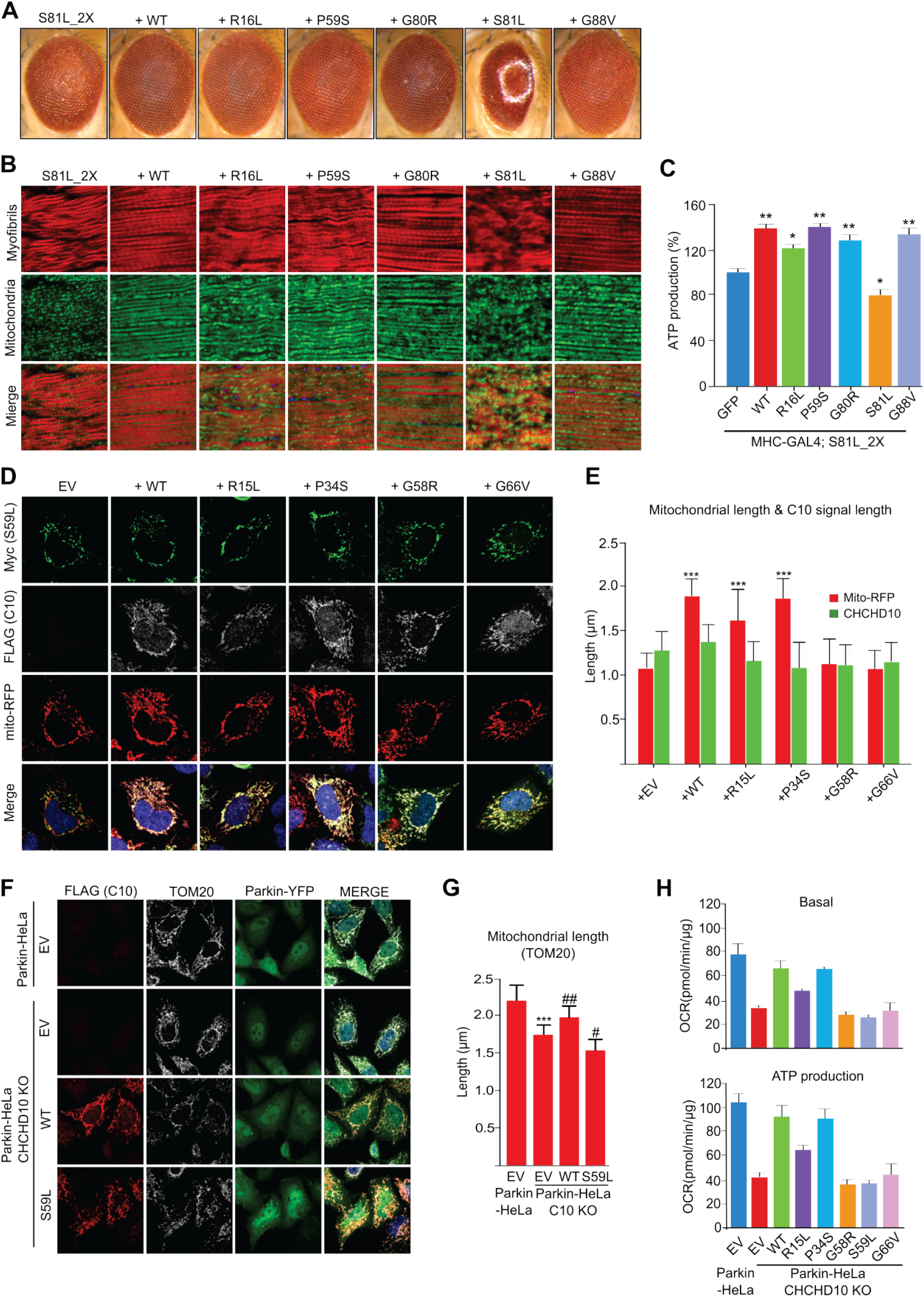
*CHCHD10^S59L^*-induced phenotypes are rescued by *CHCHD10* co-expression. (**A**) Expression of *C2C10H^WT^*, *C2C10H^R16L^*, *C2C10H^P59S^*, *C2C10H^G80R^*, and *C2C10H^G88V^* with two copies of *C2C10H^S81L^* by *GMR*-GAL4 improved *C2C10H^S81L^*-induced rough eye phenotypes. (**B**) Expression of *C2C10H^WT^*, *C2C10H^R16L^*, *C2C10H^P59S^*, *C2C10H^G80R^*, and *C2C10H ^G88V^* with two copies of *C2C10H^S81L^* in muscle tissues improved muscle architecture and abnormal mitochondrial phenotypes. (**C**) ATP levels in each of the indicated genotypes were measured as shown in Fig. 2E. Data are mean ± SE (one-way ANOVA, \**P* < 0.05, \*\**P* < 0.001; *n* = 3 replicates). (**D**) HeLa cells were co-transfected with Myc-tagged *CHCHD10^S59L^* and each FLAG-tagged variant. Empty vector (EV) was used as a control. *mTagRFP-T-Mito-7* was co-transfected to visualize mitochondrial localization of CHCHD10 proteins. Representative images of HeLa cells stained with anti-Myc (green) and anti-FLAG (gray) antibodies 24 hours after transfection. (E) Quantification of Mito-RFP and CHCHD10^S59L^-Myc signal strength. Data shown are mean ± SD (one-way ANOVA followed by Tukey posthoc comparison with EV, \*\*\**P* < 0.0001; *n* = 3 biological replicates, with ≥ 15 cells for each sample). (**F**) Representative images of *CHCHD10^KO^* HeLa^YFP-Parkin^ cells transfected with *CHCHD10^WT^* and *CHCHD10^S59L^*. Cells were immunostained with antibodies against FLAG (red, CHCHD10) and TOM20 (gray, mitochondria). (**G**) Quantification of TOM20 signal strength (mitochondria). Data shown are mean ± SD (one-way ANOVA followed by posthoc Tukey tests, \*\*\**P* < 0.0001 *vs.* EV, #*P* < 0.05, ##*P* < 0.01 *vs. CHCHD10^KO^*; *n* = 3 biological replicates, with ≥ 20 cells for each sample). (**H**) *CHCHD10^KO^* HeLa^YFP-Parkin^ cells were transfected with FLAG-tagged EV, *CHCHD10^WT^*, or each of the variants. After 24 hours, mitochondrial respiration for each transfectant was measured via Seahorse XF Cell Mito Stress tests. Data shown are mean ± SD (*n* = 3 technical replicates).

In HeLa cells, co-expression of *CHCHD10^R15L^* or *CHCHD10^P34S^* with *CHCHD10^S59L^* clearly mitigated morphologic (Fig. 3D and E) and functional defects (fig. S3A), but co-expression of the other variants did not affect *CHCHD10^S59L^*-mediated degeneration. However, the punctate staining pattern of CHCHD10^S59L^ was not altered by *CHCHD10^WT^* co-expression (Fig. 3D and E), indicating that CHCHD10^WT^ improves mitochondrial integrity by a mechanism other than restoring mutant protein mislocalization. Genetic interaction studies with various *CHCHD10* mutants in *Drosophila* revealed that all of the mutants, including an *N*-terminal deletion mutant and a hydrophobic region deletion, maintained WT activity. Only a mutant lacking a *C*-terminal CHCHD domain failed to rescue the *C2C10H^S81L^*-induced phenotype (fig. S3D), which may reflect the failure of CHCHD10^ΔCHCHD^ mitochondrial localization, as previously reported (*34*).

### CHCHD10^S59L^ is a dominant gain-of-function mutant

If *C2C10H^S81L^* is a dominant-negative mutant that suppresses C2C10H^WT^ activity, the *C2C10H* deletion mutant (*C2C10H^null^*) phenotype will not be enhanced by *C2CH10^S59L^*. Because *C2C10H^null^* flies do not exhibit an abnormal eye phenotype and are generally healthy (*35*), we hypothesized that the *C2C10H^S81L^*-induced rough eye phenotype would not occur in the *C2CH10H^null^* background. However, the rough eye phenotype was robust without *C2C10H^WT^* expression (fig. S3E). This suggests that *C2C10H^S81L^* is a dominant gain-of-function mutation that does not enhance its own activity but instead acquires a new toxic mechanism that can be mitigated by *C2C10H^WT^* co-expression.

To further validate this result in a mammalian system, we generated *CHCHD10* knockout (*CHCHD10^KO^*) HeLa cells via the CRISPR/Cas9 system (fig. S3F). Although the CHCHD10 protein was not detected with an anti-CHCHD10 antibody in the *CHCHD10^KO^* lines, mitochondrial morphology and respiratory function were not affected (fig. S3F and G). Surprisingly, *CHCHD10^KO^* HeLa cells stably expressing *YFP-Parkin* exhibited marked morphologic defects and decreased respiratory function (Fig. 3F–H and fig. S3H). These defects were recovered by transiently expressing *CHCHD10^WT^*, whereas expression of *CHCHD10^S59L^* exacerbated the morphologic and functional defects (Fig. 3F–H). Therefore, *CHCHD10^S59L^* is a dominant gain-of-function mutation. Moreover, expression of *CHCHD10^R15L^* or *CHCHD10^P34S^* recovered respiratory function (Fig. 3H), confirming that these mutants retain WT activity.

### CHCHD10 variants form insoluble aggregates

The punctate staining pattern of CHCHD10^S59L^ was not affected when CHCHD10^WT^ rescued CHCHD10^S59L^-induced mitochondrial fragmentation. Immunostaining of indirect flight muscles expressing *C2C10H^S81L^* revealed that C2C10H^S81L^ proteins formed aggregate-like structures *in vivo* (fig. S4A). The G58R, S59L, and G66V substitutions are located in the hydrophobic domain of an intrinsically disordered region in CHCHD10 (Fig. 4A). Because the hydrophobic domain and intrinsically disordered region play a role in protein folding and aggregation, we examined whether mutant CHCHD10 proteins accumulate in mitochondria as insoluble aggregates. We detected insoluble CHCHD10^S59L^ and CHCHD10^G66V^ proteins (i.e., RIPA insoluble and urea soluble) by immunoblotting (Fig. 4B and fig. S4B), which was consistent with the presence of mitochondrial punctate structures in HeLa cells containing these variants (Fig. 2A). However, we did not observe insoluble CHCHD10^G58R^, although CHCHD10^G58R^ was clearly visualized in punctate structures. Co-expression of *CHCHD10^WT^* with *CHCHD10^S59L^* did not suppress accumulation of insoluble CHCHD10^S59L^ (Fig. 4B) but recovered mitochondrial morphology and function (Fig. 4B).

**Fig. 4.**
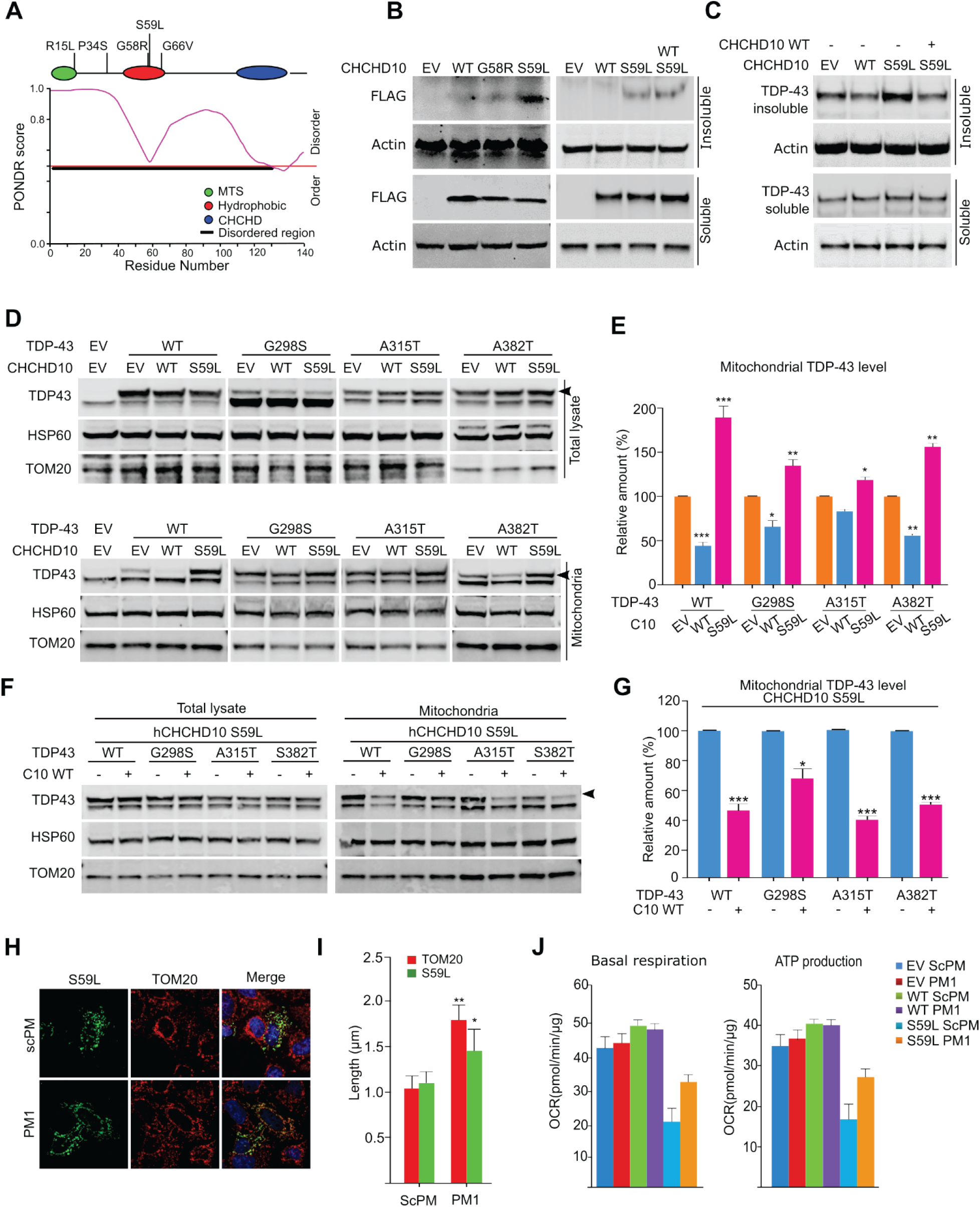
*CHCHD10^S59L^* increases TDP-43 insolubility and mitochondrial translocation. (**A**) The intrinsically disordered region of CHCHD10 was predicted by PONDR. (**B**) HeLa cells were transfected with FLAG-tagged *CHCHD10^WT^*, *CHCHD10^G58R^*, or *CHCHD10^S59L^*. After 24 hours, cells were subjected to sequential protein extraction with RIPA and urea buffers. Immunoblotting was conducted with anti-FLAG and anti-actin (loading control) antibodies. (**C**) HeLa cells transfected with FLAG-tagged *CHCHD10^WT^* or *CHCHD10*^S59L^ with *TARDBP* were subjected to sequential protein extraction with RIPA and urea buffers. Immunoblotting was performed with anti-TDP-43 or anti-actin (loading control) antibodies. (**D** and **E**) HeLa cells were co-transfected with FLAG-tagged *TARDBP^WT^* or disease-causing mutants combined with empty vector (EV), *CHCHD10^WT^*, or *CHCHD10^S59L^*. After 24 hours, mitochondria were fractionated. Immunoblotting was conducted with anti-TDP-43 (arrowhead indicates FLAG-TDP-43), anti-HSP60, and anti-TOM20 (loading control) antibodies. Data shown are mean ± SD (one-way ANOVA, \**P* < 0.05, \*\**P* < 0.01, \*\*\**P* < 0.001; *n* = 3). (**F** and **G**) HeLa cells were co-transfected with FLAG-tagged *TARDBP^WT^* or disease-causing mutants combined with EV or *CHCHD10^WT^*. After 24 hours, mitochondria were fractionated. Immunoblotting was conducted with anti-TDP-43 (arrowhead indicates FLAG-TDP-43) or anti-HSP60 and anti-TOM20 (loading control) antibodies. Data shown are mean ± SD (one-way ANOVA, \**P* < 0.05, \*\*\**P* < 0.001; *n* = 3). (**H**) HeLa cells expressing *CHCHD10^S59L^* were treated with a control peptide (scPM, 5 μM) or TDP-43 inhibitor (PM1, 5 μM). Representative images of HeLa cells stained with FLAG (green, CHCHD10^S59L^) and TOM20 (red, mitochondria) 24 hours after transfection. (**F**) Quantification of TOM20 and CHCHD10^S59L^-FLAG signal strength. Data are mean ± SD (one-way ANOVA, \**P* < 0.05, \*\**P* < 0.01; *n* = 3 biological replicates, with ≥ 20 cells in each sample). (**J**) Mitochondrial respiration was measured by Seahorse XF Cell Mito Stress tests. Cells were treated with 2 μM ScPM or PM1.

### Expression of CHCHD10^WT^ reduces TDP-43 insolubility and mitochondrial translocation

Mutations in *TARDBP* cause ALS and FTD, and cytoplasmic TDP-43 aggregates are a hallmark in most patients with ALS and/or FTD (*36, 37*). Therefore, we examined whether *CHCHD10^S59L^* genetically interacts with mutant *TARDBP* in *Drosophila*. Co-expression of disease-causing *TARDBP^M337V^* with *CHCHD10^S81L^* in *Drosophila* eyes with *GMR*-GAL4 caused synergistic lethality (due to leaky expression in other tissues). Replacing the entire coding region of *Drosophila TBPH* with human *TARDBP^M337V^* cDNA (*38*) enhanced the *CHCHD10^S59L^*-induced rough eye phenotype (data not shown). We next examined insoluble TDP-43 levels after *CHCHD10^WT^* and *CHCHD10^S59L^* transfection in HeLa cells. *CHCHD10^WT^* expression decreased accumulation of insoluble TDP-43, whereas *CHCHD10^S59L^* expression increased insoluble TDP-43 (Fig. 4C). Co-expression of *CHCHD10^WT^* and *CHCHD10^S59L^* suppressed *CHCHD10^S59L^*-induced insoluble TDP-43 (Fig. 4C).

We then determined whether *CHCHD10^WT^* and *CHCHD10^S59L^* expression affects mitochondrial translocation of TDP-43. Despite similar expression levels in whole-cell lysates, exogenously expressed *TARDBP^WT^* and three pathogenic mutants (G298S, A315T, and A382T) showed increased mitochondrial distribution in *CHCHD10^S59L^*-expressing cells over that of empty vector–transfected cells. We observed decreased mitochondrial distribution of TDP-43^WT^ and its mutants in *CHCHD10^WT^*-expressing cells (Fig. 4D and E). Furthermore, co-expression of *CHCHD10^WT^* and *CHCHD10^S59L^* reduced mitochondrial translocation of all TDP-43 variants (Fig. 4F and G). Immunofluorescence staining confirmed the increased mitochondrial localization of TDP-43 in *CHCHD10^S59L^*-expressing cells (fig. S4C). To test whether inhibition of TDP-43 mitochondrial translocation recovers *CHCHD10^S59L^*-induced mitochondrial morphologic and functional defects, we treated *CHCHD10^S59L^*-transfected cells with the PM1 peptide inhibitor of TDP-43 mitochondrial translocation (*25*). Notably, the morphologic and functional defects caused by *CHCHD10^S59L^* were ameliorated by PM1 (Fig. 4H–J and fig. S4D), suggesting that increased TDP-43 insoluble aggregation and mitochondrial translocation are critical to *CHCHD10^S59L^*-induced toxicity.

### PINK1/parkin mediates dominant toxicity in the C2C10H^S81L^ Drosophila model

Because only *CHCHD10^S59L^* expression caused consistent dominant toxicity in both *Drosophila* and HeLa cells, we investigated the mechanism of action of *CHCHD10^S59L^*-mediated toxicity. Mitochondrial fragmentation is a major phenotype caused by expression of *CHCHD10^S59L^*. Therefore, we hypothesized that the genes involved in mitochondrial dynamics or mitochondrial quality control are effectors of *CHCHD10^S59L^*-driven mitochondrial pathogenesis. We performed genetic interaction studies by using the *Drosophila C2C10H^S81L^* eye model with various RNAi, classical deficiency, and duplication lines (supplemental table 1). The strongest dominant suppressor we identified was PINK1. Down-regulation of *Pink1* by RNAi rescued the rough eye phenotype produced by two copies of *C2C10H^S81L^*, whereas *Pink1* overexpression enhanced the rough eye phenotype (Fig. 5A). RNAi-mediated depletion of *park* (i.e., the *Drosophila* gene encoding parkin), a downstream partner of PINK1 in mitochondrial quality control, also marginally rescued the rough eye phenotype (Fig. 5A).

**Fig. 5.**
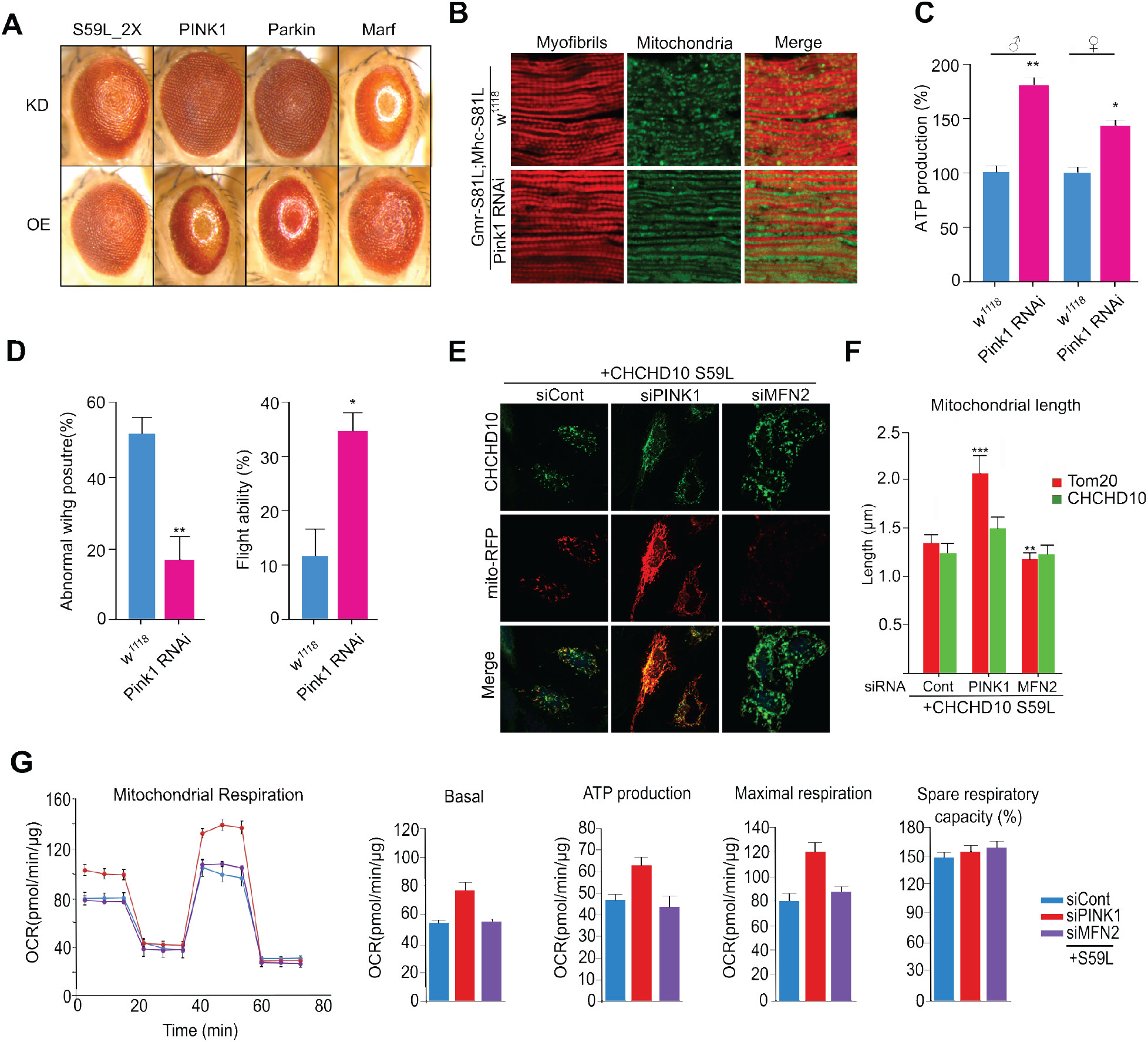
*CHCHD10^S59L^*-induced degeneration is rescued by *Pink1* downregulation. (**A**) RNAi-mediated knockdown of *Pink1* or *Park* rescued *C2C10H^S81L^*-induced eye phenotypes, and overexpression of *Pink1* or *Park* exacerbated the *C2C10H^S81L^*-induced eye phenotype. (**B**) RNAi-mediated knockdown of *Pink1* in muscles rescued *C2C10H^S81L^*-induced muscle degeneration and mitochondrial defects. (**C**) ATP levels in thoraxes from 10-day-old flies were measured. Data are mean ± SD (one-way ANOVA, \**P* < 0.05, \*\**P* < 0.01; *n* = 3 replicates) (**D**) Down-regulation of *Pink1* increased ATP production and mitigated abnormal wing postures and locomotor defects assessed by flight ability. Data are mean ± SD (one-way ANOVA, \**P* < 0.05, \*\**P* < 0.01; *n* = 3 replicates) (**E**) HeLa cells were transfected with siRNAs targeting *PINK1* or *MFN2*. After 18 hours, the cells were transfected with *CHCHD10^S59L^* and *Mito-RFP*. Representative images of transfected HeLa cells immunostained with antibodies against FLAG. (**F**) Quantification of Mito-RFP and CHCHD10^S59L^-FLAG signal strength. Data shown are mean ± SD (one-way ANOVA followed by Tukey posthoc comparison with EV; \*\*\**P* < 0.001; *n* = 3, with ≥ 20 cells measured for each sample). (**G**) Mitochondrial respiration was measured by Seahorse XF Cell Mito Stress tests 24 hours after *CHCHD10* transfection. Data shown are mean ± SD (*n* = 3 technical replicates).

We also tested several other genes that are parallel or downstream of the PINK1/parkin pathway, including *mul1*, *ari1*, *march5*, *Drp1*, *marf*, and *TER94* (supplemental table 1). Only overexpression of *marf*, a *Drosophila* ortholog of *MFN* (i.e., mitofusin), showed slight suppressive effects with consistent RNAi-mediated enhancement (Fig. 5A). This indicates that Marf may be a downstream effector of the PINK1/parkin pathway during *CHCHD10^S59L^*-mediated pathogenesis. However, this does not exclude other genes from playing a role in *C2CH10^S81L^*-mediated pathogenesis. RNAi-mediated depletion of *Pink1* also recovered *C2C10H^S81L^*-dependent indirect flight muscle degeneration (Fig. 5B). *Pink1* knockdown rescued sarcomere structure, extended mitochondrial length, and increased ATP production (Fig. 5B and C), improving *C2C10H^S81L^*-induced defects, including abnormal wing posture and flight ability (Fig. 5D).

### PINK1/parkin mediates dominant toxicity in HeLa cells

RNAi-mediated knockdown of *PINK1* remarkably rescued *CHCHD10^S59L^*-induced mitochondrial morphologic and functional defects in HeLa cells (Fig. 5E–G and fig. S5A). Furthermore, expression of *CHCHD10^S59L^* in *PINK1^KO^* HeLa cells did not cause mitochondrial fragmentation (fig. S5B). Although *MFN2* knockdown did not affect *CHCHD10^S59L^*-induced mitochondrial defects in HeLa cells (Fig. 5E–G), simultaneous knockdown of *MFN1* and *MFN2* enhanced the mitochondrial defects (fig. S5C and D). Overexpression of *MFN2-YFP* was uninterpretable because of its strong mitochondrial clustering activity (*39*), although it marginally improved respiratory function (fig. S5E and F).

In *Drosophila*, both *Drp1* knockdown and overexpression yielded detrimental effects (supplemental table 1). However, RNAi-mediated depletion of *DRP1* in HeLa cells successfully reversed *CHCHD10^S59L^*-induced mitochondrial fragmentation and improved respiratory function (fig. S5G–I). Consistently, *CHCHD10^S59L^*-mediated respiratory defects were slightly enhanced by *DRP1-YFP* co-expression (fig. S5F). Indeed, CHCHD10^S59L^ aggregates were nearly absent with *DRP1* knockdown, in contrast with persistent aggregates with *PINK1* knockdown. We speculated that pre-elongated mitochondria with *DRP1* knockdown reduces local concentration of CHCHD10^S59L^ proteins, thereby preventing aggregate formation. Although modulating mitochondrial dynamics may be beneficial in reducing *CHCHD10^S59L^*-induced toxicity, modulating PINK1, parkin, or MFN is less detrimental than directly modulating DRP1.

### Parkin-mediated mitophagy induces toxicity

PINK1 accumulates in mitochondria upon mitochondrial stress or damage (*40*) and recruits parkin by phosphorylating ubiquitin and other substrates, including parkin, resulting in MFN1/2 degradation and mitophagy to remove damaged mitochondria (*41–44*). The suppressive effects of *PINK1* and *PRKN* knockdown suggest that CHCHD10^S59L^ induces PINK1 accumulation in mitochondria, activating the PINK1/parkin pathway. PINK1-YFP accumulated in the mitochondria of *CHCHD10^S59L^*-expressing HeLa cells, in contrast with that of *CHCHD10^WT^*-overexpression (Fig. 6A and C). Because *PRKN* expression is deficient in HeLa cells, we used stable expression of *YFP-Parkin* (HeLa^YFP-Parkin^) to study parkin-mediated mitophagy. When *CHCHD10^S59L^* was transiently expressed in HeLa^YFP-Parkin^ cells, YFP-Parkin was also recruited to mitochondria (Fig. 6B and D), clearly demonstrating that CHCHD10^S59L^ induces PINK1 stabilization, accumulation in mitochondria, and subsequent parkin recruitment. This suggests that PINK1/parkin-mediated mitophagy is toxic in this system. Indeed, enhancing autophagy by *Atg1* expression in the *C2C10H^S81L^* eye model was synergistically lethal (supplemental table 1), although beneficial effects of *Atg1* overexpression are reported in *Drosophila* (*45, 46*). However, strong mitochondrial abnormalities in *PRKN*-deficient HeLa cells indicate the presence of parkin-independent toxic mechanisms. We therefore hypothesized that PINK1-mediated, parkin-independent basal mitophagy is highly activated in HeLa cells and that dysregulated basal mitophagy is amplified in the presence of parkin. To test this hypothesis, we assessed LC3 conversion and accumulation in mitochondria to examine whether autophagosome formation is activated by *CHCHD10^S59L^* expression in HeLa cells.

**Fig. 6.**
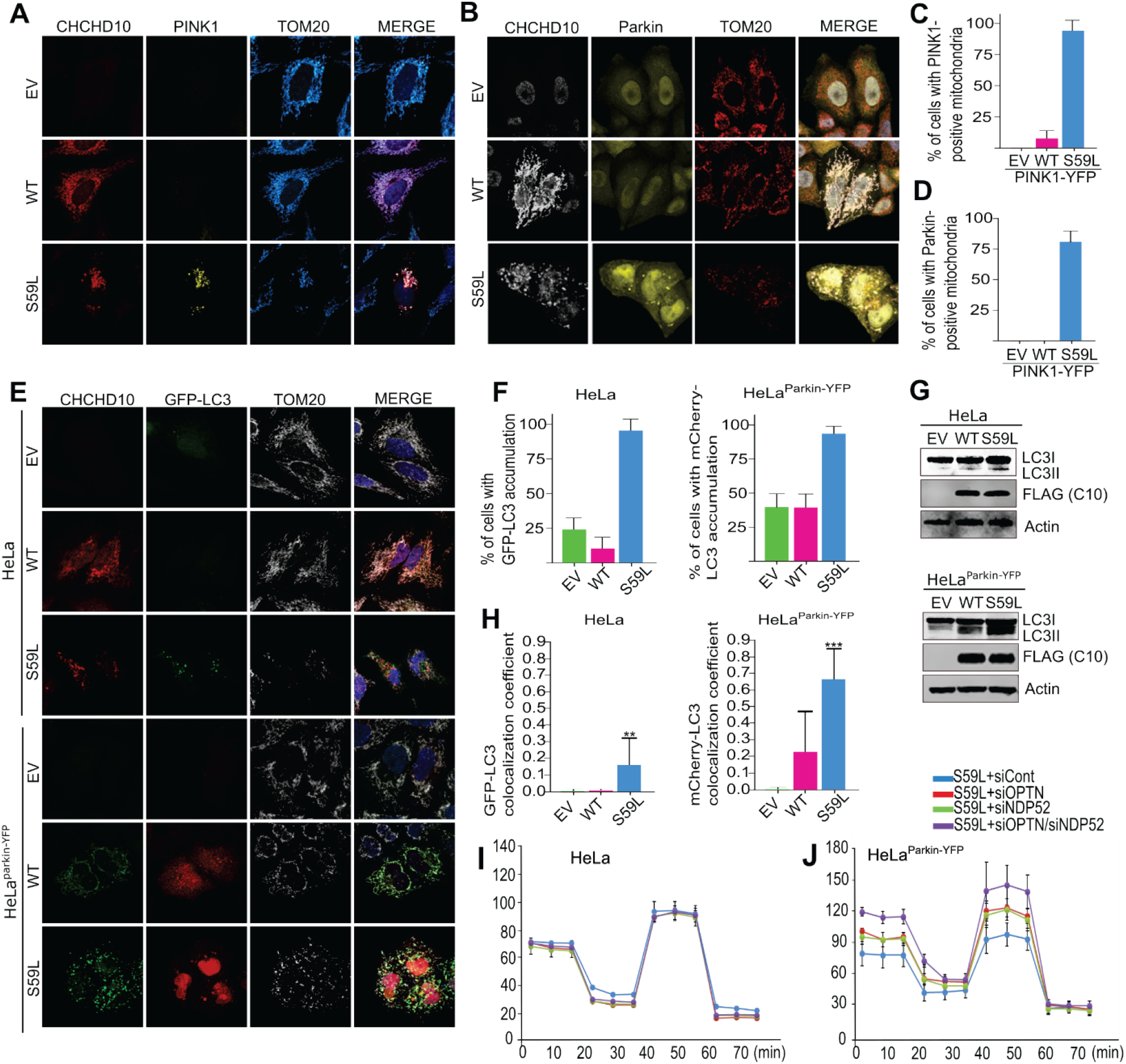
*CHCHD10^S59L^*-induced toxicity is mediated by parkin-mediated mitophagy. (**A**) HeLa cells and (**B**) HeLa^YFP-Parkin^ cells were transfected with FLAG-tagged *CHCHD10^S59L^* and *PINK1-YFP*. The transfected HeLa cells were immunostained with antibodies against FLAG and TOM20 24 hours after transfection. (**C** and **D**) Percentage of HeLa cells showing (C) PINK1-YFP and (D) YFP-Parkin accumulation in mitochondria; *n* = 3, with ≥ 15 cells per condition. (**E**) HeLa cells and HeLa^YFP-Parkin^ cells were co-transfected with FLAG-tagged *CHCHD10^S59L^* and *GFP-LC3* or *mCherry-LC3*, respectively. Representative images of transfected cells immunostained with antibodies against FLAG and TOM20. (**F**) Percentage of cells showing GFP-or mCherry-LC3 accumulation (mean ± SD; \*\*\**P* < 0.001; *n* = 3, with ≥ 10 cells for each group). (**G**) Immunoblot of LC3 and CHCHD10^S59L^ expression. (**H**) GFP-LC3 and mCherry-LC3 co-localization with TOM20 in HeLa cells and HeLa^YFP-Parkin^ cells. (**I**) HeLa cells and (**J**) HeLa^YFP-Parkin^ cells were transfected with siRNAs targeting *NDP52* or/and *OPTN*. At 24 hours after siRNA transfection, mitochondrial respiration was measured by Seahorse XF Cell Mito Stress tests (*n* = 3 technical replicates).

In both HeLa and HeLa^YFP-Parkin^ cells, *CHCHD10^S59L^* expression increased LC3 accumulation (Fig. 6E and F) and LC3 conversion (Fig. 6G). However, only small portions of LC3 accumulation co-localized with mitochondria, as compared with strong mitochondrial co-localization of LC3 accumulation in HeLa^YFP-Parkin^ cells (Fig. 6E and H). Co-staining of lysosomes and mitochondria also revealed limited lysosomal staining in the mitochondria of *CHCHD10^S59L^*-expressing HeLa cells (fig. S6A). Therefore, expression of *CHCHD10^S59L^* induces general nonspecific autophagy without parkin, whereas mitophagy induction is a major phenomenon in HeLa^YFP-Parkin^ cells. Consequently, RNAi-mediated depletion of the basal mitophagy receptors *NDP52* and optineurin (*OPTN*) had no effect on *CHCHD10^S59L^*-mediated toxicity in HeLa cells but increased respiratory activity in HeLa^YFP-Parkin^ cells (Fig. 6I and J and fig. S6B–D). Therefore, the PINK1/parkin pathway is a major toxicogenic pathway, but PINK1 downstream factors mediate this toxicity independent of parkin.

### PINK1 activation does not disrupt mitochondrial membrane potential or activate the UPR

To investigate how CHCHD10^S59L^ induces PINK1 accumulation, we examined whether expression of *CHCHD10^S59L^* disrupts mitochondrial membrane potential at a single-cell level. We used a split GFP strategy because of possibly mislocalized *C*-terminal GFP-tagged CHCHD10 (*34*). The combination of GFP11-tagged CHCHD10^S59L^ with mitochondrial-localized GFP1–10 revealed that CHCHD10^S59L^ does not disrupt mitochondrial membrane potential, although it induced mitochondrial fragmentation (fig. S7A). The unfolded protein response (UPR) in mitochondria (UPRmt) induces PINK1 accumulation without disrupting mitochondrial membrane potential (*40*). We examined the protein levels of two UPR markers, CHOP and HSP60, which are increased upon UPRmt (*47*). Although UPR-inducing mutant ornithine transcarbamylase (*ΔOTC*) increased CHOP and HSP60 levels, these proteins were not increased by *CHCHD10^S59L^* (fig. S7B). A genetic interaction of increased UPRmt with *ΔOTC* expression did not occur in *Drosophila* expressing *C2C10H^S81L^* (fig. S7C). Therefore, UPRmt is not a major cause of *CHCHD10^S59L^*-induced degeneration and PINK1 accumulation.

### Modulating PINK1 downstream pathways mitigates CHCHD10^S59^-induced toxicity

To further define the parkin-independent downstream pathway of PINK1, we tested whether the known downstream components ND42, sicily, Miro, and mitofilin modulate the *C2CH10^S81L^*-induced rough eye phenotype (supplemental table 1). Only co-expression of PINK1 phosphorylation-null *Mitofilin* (*48*) showed mild but obvious rescue of the *C2C10H^S81L^*-induced rough eye phenotype (fig. S7D), whereas RNAi-mediated knockdown of *Mitofilin* enhanced this phenotype (supplemental table 1). Similarly, co-expression of PINK1 phosphorylation-null *MFN2^S378A^* (*49*) with *CHCHD10^S59L^* in HeLa cells attenuated the abnormal mitochondrial phenotype (Fig. 7A). These data suggest that fine modulation of the downstream events caused by PINK1-mediated phosphorylation is an effective therapeutic strategy. Indeed, recently developed MFN2 agonists rescued *CHCHD10^S59L^*-induced morphologic and functional mitochondrial defects (Fig. 7B and C and fig. S7E). We also observed moderately elongated mitochondria in the indirect flight muscles of *C2C10H^S81L^*-expressing flies reared on two MFN2 agonists (Fig. 7D), which also had increased ATP production in flies expressing *C2C10H^S81L^* (Fig. 7E) and in *C9orf72*-expressing flies with expanded GGGGCC repeats (fig. S7F) (*50*).

**Fig. 7.**
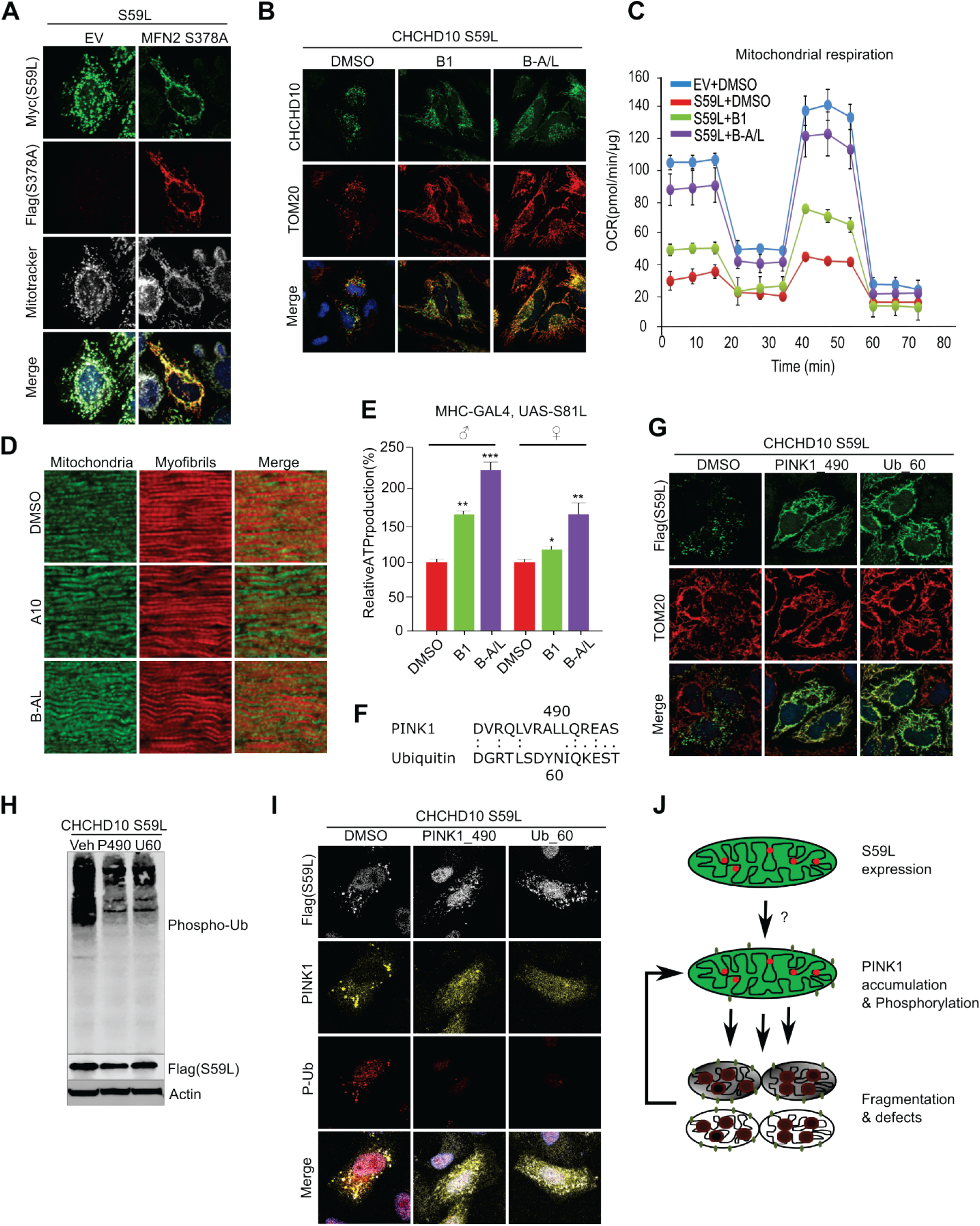
*CHCHD10^S59L^*-induced toxicity is mitigated by inhibiting PINK1 activity and its target molecules. (**A**) HeLa cells were transfected with Myc-tagged *CHCHD10^S59L^* and FLAG-tagged *MFN2^S378A^*. Mitochondria were stained with mitotracker (deep red FM) and antibodies against Myc and FLAG 24 hours after transfection. (**B**) HeLa cells were transfected with FLAG-tagged *CHCHD10^S59L^* and treated with MFN2 agonists B1 (50 nM) and B-A/L (5 nM) for 24 hours. Cells were immunostained with antibodies against FLAG and TOM20. (**C**) Mitochondrial respiration was measured by Seahorse XF Cell Mito Stress tests. Graphs show the basal, maximal, and spare respiratory capacity (%) and ATP production. (**D**) Indirect flight muscles from 10-day-old adult flies fed with B1 and B-A/L (each 10 μM) were stained with streptavidin– Alexa Fluor 488 and phalloidin–Alexa Fluor 594. DMSO was used as a vehicle. (**E**) ATP levels in thoraxes from the indicated genotypes (aged 10 days) were measured. Data are mean ± SE (one-way ANOVA, \**P* < 0.05, \*\**P* < 0.01, and \*\*\**P* < 0.001; *n* = 3). (**F**) Peptide sequences of two PINK1 inhibitors. (**G**) HeLa cells were treated with PINK1_490 (0.15 µg/ml) or Ub_60 (0.15 µg/ml) and transfected with FLAG-tagged *CHCHD10^S59L^*. DMSO was used as a vehicle. Cells were immunostained with antibodies against FLAG and TOM20 24 hours after *CHCHD10^S59L^* transfection. (**H**) HeLa cells treated with peptide inhibitors and transfected with FLAG-tagged *CHCHD10^S59L^* were analyzed with an anti–phospho-ubiquitin antibody. (**I**) HeLa cells were treated with peptide inhibitors followed by transfection of FLAG-tagged *CHCHD10^S59L^* and *PINK1-YFP*. The cells were immunostained with antibodies against FLAG and TOM20. (**J**) A schematic depicting the positive feedback mechanism of *CHCHD10^S59L^*-mediated toxicity through PINK1 accumulation.

### Inhibition of PINK1 activity mitigates CHCHD10^S59^-induced toxicity

We hypothesized that inhibiting PINK1 kinase activity is a more effective and robust therapeutic strategy than regulating PINK1 expression levels or its substrates. Because of a lack of small-molecule inhibitors of PINK1, we used LALIGN to generate peptide sequences that may act as pseudo-substrate inhibitors of PINK1 and ubiquitin (Fig. 7F) (*51, 52*). Although treating HeLa cells with two peptides via a protein delivery reagent did not produce *CHCHD10^S59L^*-mediated mitochondrial fragmentation in both pretreated and post-treated experiments (Fig. 7G and fig. S7G), ubiquitin phosphorylation was reduced (Fig. 7H and I). Peptide treatment also reduced PINK1 accumulation and phospho-ubiquitin staining (Fig. 7I), suggesting that a positive feedback mechanism amplifies *CHCHD10^S59L^*-mediated toxicity via PINK1 accumulation (Fig. 7J).

### PINK1 knockdown does not affect TDP-43, but CHCHD10^WT^ prevents PINK1 accumulation

To determine whether the two axes of *CHCHD10^S59L^*-induced toxicity exhibit crosstalk, we examined TDP-43 insolubility and mitochondrial localization after RNAi-mediated *PINK1* knockdown in *CHCHD10^S59L^*-transfected HeLa cells. Although *PINK1* knockdown strongly rescued the morphologic and functional defects in *Drosophila* and HeLa cells, insoluble TDP-43 levels and mitochondrial localization of TDP-43^WT^ and TDP-43^A382T^ were unaffected in HeLa cells (Fig. 8A and B). Additionally, insoluble CHCHD10^S59L^ levels were not changed by *PINK1* knockdown (Fig. 8B), suggesting that PINK1 accumulation in response to *CHCHD10^S59^*^L^ expression is parallel or downstream (or both) of the TDP-43 pathway. *CHCHD10^WT^* co-expression with *CHCHD10^S59L^* reduced PINK1 accumulation (Fig. 8C). *CHCHD10^WT^* expression also reduced PINK1 and parkin accumulation caused by mild CCCP treatment in *PINK1*-expressing HeLa and HeLa^YFP-Parkin^ cells, respectively (Fig. 8D and fig. S8). Therefore, *CHCHD10^WT^* protected mitochondria not only through TDP-43 but also by directly preventing PINK1 accumulation. *CHCHD10^WT^* exerted a protective effect through both TDP43 and PINK1 without modulating CHCHD10^S59L^ insolubility (Fig. 8E), suggesting that augmenting *CHCHD10* expression or activity is also a promising therapeutic strategy.

**Fig. 8.**
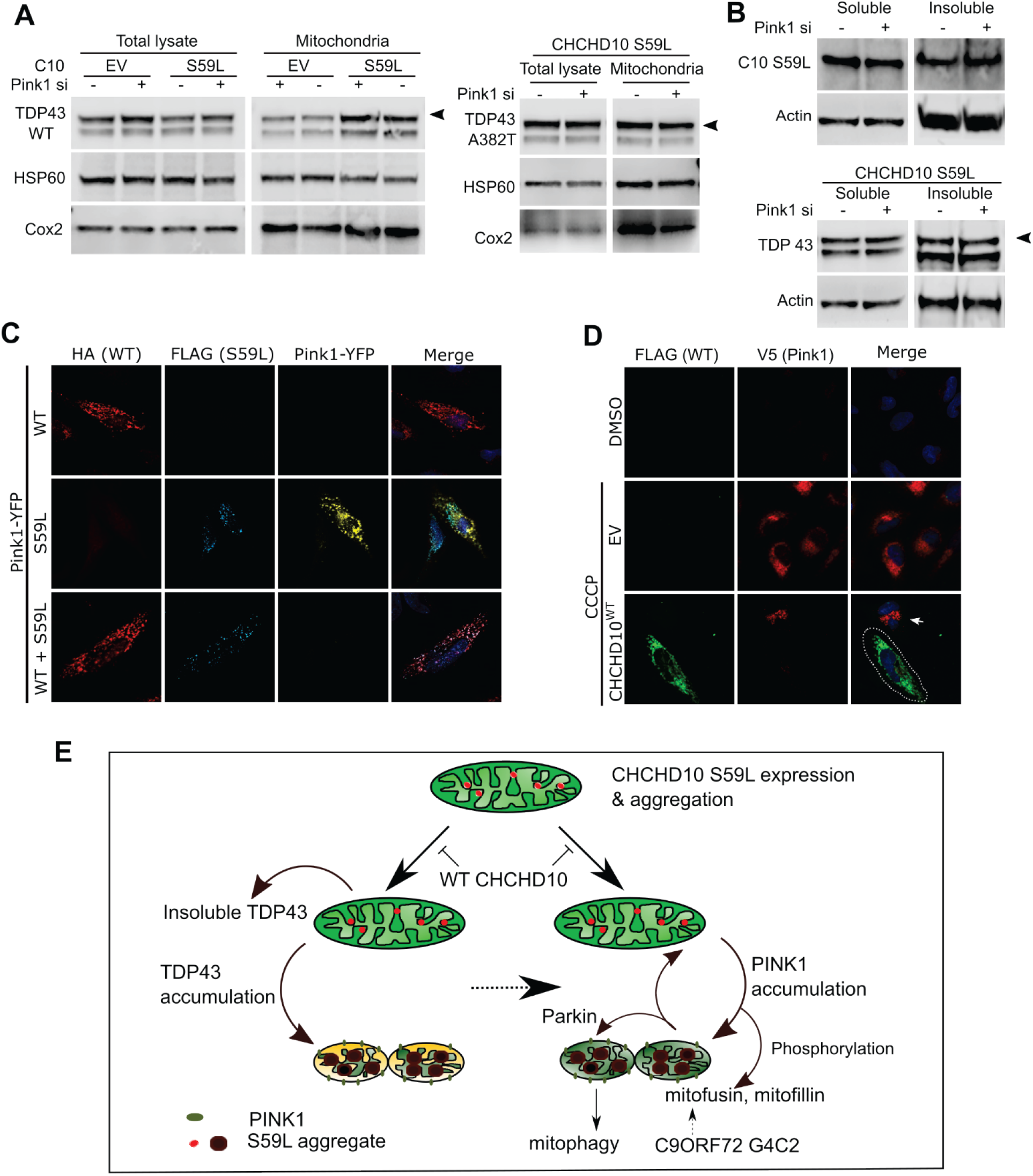
Dominant toxicity of *CHCHD10^S59L^* is mediated independently by TDP-43 and PINK1 signaling. (**A**) HeLa cells transfected with *PINK1* siRNA were co-transfected with empty vector (EV) or FLAG-tagged *CHCHD10^S59L^* and FLAG-tagged *TARDBP^WT^* or *TARDBP^A382T^*. Fractionated mitochondria were analyzed with anti-TDP-43 (arrowhead indicates FLAG-TDP-43) or anti-HSP60 and anti-Cox2 (loading control) antibodies. (**B**) HeLa cells transfected with *PINK1* siRNA were co-transfected with FLAG-tagged *CHCHD10^S59L^* (upper panel) or FLAG-tagged *CHCHD10^S59L^* and *TARDBP* (lower panel). RIPA soluble and insoluble fractions were analyzed with anti-FLAG, anti-TDP43, or anti-actin (loading control) antibodies. (**C**) HeLa cells transfected with *PINK1-YFP*, FLAG-tagged *CHCHD10^S59L^*, and HA-tagged *CHCHD10^WT^* or EV were visualized with anti-FLAG (blue) and anti-HA (red) antibodies and YFP (yellow) after 24 hours of transfection. (**D**) HeLa^PINK1-V5-His^ cells transfected with EV or *CHCHD10^WT^* were treated with CCCP (10 μM) for 6 hours. The cells were analyzed with anti-FLAG (green) and anti-V5 (red) antibodies to visualize CHCHD10 and PINK1, respectively. Arrow indicates PINK1 accumulated in a nontransfected cell neighboring a *CHCHD10*-transfected cell (white dashed line). (**E**) Graphical summary of *CHCHD10^S59L^*-induced toxicity.

## Discussion

Since identifying the S59L substitution in CHCHD10, additional variants have been identified and suggested as pathogenic mediators of ALS-FTD, SMAJ, and mitochondrial myopathy. Despite efforts to elucidate their pathogenic mechanisms, many controversial findings suggest that mutations in *CHCHD10* do not share a common disease-causing mechanism (*1, 11, 12, 19–22, 25*). Our results also mirror these current controversial findings. We found that only *CHCHD10^S59L^* induced consistent dominant toxicity in both *Drosophila* and HeLa cells. *CHCHD10^P34S^* was not pathogenic in our experimental systems and was equivalent to *CHCHD10^WT^* in every aspect we tested. Although *CHCHD10^R15L^* haploinsufficiency is a proposed mechanism and *C2C10H^R16L^* did not cause any detectable defects in *Drosophila*, overexpression of *CHCHD10^R15L^* causes mild defects in mammalian cells (*19, 21*). However, CHCHD10^R15L^ retained WT-like activity that mitigated *CHCHD10^S59L^*-induced toxicity in *Drosophila*, HeLa cells, and in *CHCHD10^KO^* cell lines. Because only mutant transcript levels are decreased in patient fibroblasts carrying *CHCHD10^R15L^*, augmenting expression of *CHCHD10^WT^* may be beneficial in this case.

Our findings with *CHCHD10^G58R^* and *CHCHD10^G66V^* are more difficult to interpret because of their disparate results in *Drosophila* and HeLa cells. Although similar steady-state expression levels occurred in *Drosophila* and HeLa cells, *CHCHD10^G58R^* and *CHCHD10^G66V^* expression is considerably reduced when compared with that of *CHCHD10^WT^* or *CHCHD10^S59L^* (*34*). Therefore, we speculate that *C2C10H^G80R^* and *C2C10H^G88V^* expression levels in *Drosophila* do not reach a threshold to cause a visible phenotype but can rescue *CHCHD10^S81L^*-induced degeneration to varying degrees. In contrast, *CHCHD10^G58R^* and *CHCHD10^G66V^* expression in HeLa cells was acute, robust, and caused aggregation and dominant toxicity.

Consistent with studies demonstrating a dominant mechanism of *CHCHD10^S59L^* in *CHCHD10^KO^* mice and *CHCHD10^S59L^* knockin mice (*53, 54*), our findings do not support simple loss-of-function or haploinsufficiency mechanisms. Woo *et al*. proposed a dominant-negative mechanism of *CHCHD10^S59L^* in *C. elegans* and mammalian systems (*19*). Although their observations are similar to ours our data support a dominant gain-of-toxicity mechanism for *CHCHD10^S59L^*. Although these two mechanisms are not necessarily mutually exclusive, several aspects of our findings primarily support a dominant gain-of-toxicity mechanism: (1) Eye phenotypes driven by *C2C10H* mutations did not differ between the *C2C10H^null^* or *C2C10H^WT^* background, indicating that the mutations exert a dominant gain of toxicity rather than suppressing WT activity. (2) Genetic modifiers modulating the dominant gain of toxicity in *Drosophila* were recapitulated in mammalian cells. (3) Parkin accumulation did not occur in *CHCHD10^KO^* HeLa^YFP-Parkin^ cells but did occur with *CHCHD10^S59L^* expression. (4) CHCHD10^WT^ did not accumulate in CHCHD10^S59L^ aggregate-like structures. (5) *CHCHD10^WT^* overexpression did not affect CHCHD10^S59L^ aggregate formation and insolubility. Therefore, CHCHD10 plays a protective role in mitochondria that is also protective for *CHCHD10^S59L^*-driven dominant toxicity, which occurred independently of disrupting WT activity.

We cannot rule out the contribution of loss-of-function– or dominant-negative–like effects, and some effect of reduced WT activity most likely occurs in disease pathogenesis. Our data suggest that the dominant toxicity of *CHCHD10^S59L^* can be mitigated by co-expressing similar levels of *CHCHD10^WT^*. Therefore, it is possible that young patients carrying a heterozygous *CHCHD10^S59L^* mutation do not experience severe symptoms because WT activity blocks mutant toxicity. However, reduced WT function or a change in the ratio of WT to mutant expression may trigger disease symptoms in older patients.

Our efforts to define the downstream pathways of dominant *CHCHD10^S59L^*-mediated toxicity yielded two axes and multiple molecular targets that can be therapeutically modulated. Mitochondrial translocation of TDP-43 is a toxicity-generating mechanism in *CHCHD10^S59L^*-expressing cells. An association between TDP-43 and CHCHD10 was previously proposed (*19*), and peptide inhibition of TDP-43 mitochondrial translocation mitigated the *CHCHD10^S59L^*-induced phenotype. The effects of *CHCHD10^WT^* on TDP-43 insolubility and translocation generally support that TDP-43 is a key effector generating mitochondrial toxicity in ALS-FTD.

We identified *PINK1* and *PRKN* as strong genetic modifiers of *C2C10H^S81L^*-mediated toxicity. PINK1/parkin-mediated pathways are generally protective for cells by removing damaged mitochondria. Nevertheless, *CHCHD10^S59L^* expression induced PINK1 stabilization in mitochondria without strong disruption of mitochondrial membrane potential, and genetic/pharmacologic inhibition of PINK1 mitigated *CHCHD10^S59L^*-induced toxicity. Reducing PINK1 or parkin-mediated pathways are beneficial in *in vivo* disease models of *SOD1*, *FUS*, and *TARDBP* mutations (*55–57*). We demonstrated that MFN2 agonists enhanced ATP production in flies expressing *C9ORF72* with GGGGCC repeats. Therefore, small molecules known to regulate this pathway, including MFN2 agonists, may have therapeutic potential for this subtype of ALS-FTD.

Two downstream phosphorylation substrates of PINK1, mitofusin and mitofilin, mediated *CHCHD10^S59L^*-induced toxicity. Although fusion activity in *CHCHD10^S59L^*-expressing cells is not altered (*1*), PINK1 accumulation and subsequent inactivation of mitofusin by phosphorylation is responsible for fragmented mitochondria and the respiratory defects caused by *CHCHD10^S59L^*. An MFN2 agonist developed for CMT2 was effective for *CHCHD10^S59L^*-mediated toxicity and may also be effective for *CHCHD10^G66V^*-mediated SMAJ or CMT2. Overexpression of PINK1 phosphorylation-null *Mitofilin* also rescued *C2C10H^S81L^*-mediated toxicity, suggesting that deformation of MICOS was not based on a direct interaction but through phosphorylation by PINK1. Therefore, the degree of PINK1 stabilization may correspond with the phenotypic severity of mutant *CHCHD10*.

Augmenting CHCHD10^WT^ activity may be a promising therapeutic strategy, regardless of specific *CHCHD10* mutations. *CHCHD10^WT^* expression is increased in response to various stresses (*58*). We and others observed that *CHCHD10^WT^* expression not only rescued mutant phenotypes but also increased mitochondrial length and respiratory activity when it wais expressed alone in *Drosophila* or HeLa cells. Our findings support that pharmacologic or epigenetic augmentation of *CHCHD10* expression may benefit patients with ALS-FTD or other degenerative diseases with similar mitochondrial defects.

## Materials and Methods

### Study design

The aim of this study was to establish a *CHCHD10* mutant-induced ALS-FTD *Drosophila* model and investigate its disease mechanism to identify therapeutic targets. To this end, we first generated *Drosophila* expressing *CHCHD10* mutant variants and elucidated their toxicity. We then validated these results in mammalian cells. We investigated *CHCHD10^S59L^*-induced cytotoxicity and determined its effectors via genetic and pharmacologic methods. We examined mutant-induced morphologic and functional defects of mitochondria in *Drosophila* and HeLa cells. To delineate the *CHCHD10^S59L^*-induced degenerative pathway, we performed targeted RNAi screening in *C2C10H^S81L^*-expressing flies and identified genetic modifiers, which we validated in HeLa cells by genetic and/or pharmacologic methods. All flies for each experiment were collected under synchronized conditions, such as age, date, population, and repetition times. The *n* for each experiment is indicated in the figure legends. All experiments were blinded and randomized for outcome assessments. No outliers were excluded in the study. Detailed sample sizes, numbers of replicates, and inclusion/exclusion criteria are indicated in the figure legends or Materials and Methods.

### Image analysis and statistical analysis

Mitochondrial length was measured by using the MiNA tool set combined with ImageJ software, as previously described (*59*). Statistical analysis was performed with Prism5 (GraphPad) software. Statistical significance was determined with Student *t* tests and one-way or two-way analysis of variance (ANOVA), depending on the number of dependent variables compared. One-or two-way ANOVAs were followed by posthoc Tukey tests for significance. *P* values ≤ 0.05 were considered statistically significant.

## Acknowledgments

We thank the Bloomington Drosophila Stock Center and the VDRC Stock Center for fly lines, the Research Instrumentation Lab at the University of Minnesota Duluth for assistance. We also thank undergraduate volunteers Sierra Skudlarek, Austin Kurtz, Anna Huy, and Maddie Chalmers for helping with *Drosophila* maintenance and plasmid preparation.

## Funding

This work was supported by grants from the Muscular Dystrophy Association, the Wallin Neuroscience Discovery Fund, and an Engebretson Drug Design and Discovery Grant to N.C.K.

## Author contributions

M.B., Y.-J.C., and N.C.K. conceived the project and performed experiments. M.B., Y.-J.C., and N.C.K. analyzed data and wrote the manuscript. G.W.D. and J.P.T. provided guidance and helpful insights to design the experiments and analyze data.

## Competing interests

G.W.D. is the scientific founder of Mitochondria in Motion, Inc., which has licensed technology from Washington University St. Louis and is pursuing commercializing treatments for neurodegenerative diseases, including small-molecule mitofusin agonists.

## Data and materials availability

All data and materials are available upon request. Mitofusin agonists should be specifically requested from G.W.D.

## Supplementary Materials

### Materials and Methods

#### Generation of Drosophila lines

To generate transgenic *Drosophila* lines carrying *C2C10H*, codon-optimized *CHCHD10*, and their variants, all cDNAs were synthesized with or without a *C*-terminal FLAG tag and cloned into pUAST-*attB* by GenScript. Transgenic *Drosophila* lines were generated by BestGene with standard ΦC31 integrase-mediated transgenesis into the *attP2* site on chromosome 3 (locus 68A4), the *attP40* site on chromosome 2 (locus 25C6), or the *VK27* site on chromosome 3 (locus 89E11).

#### Drosophila genetics

Fly cultures and crosses were performed on standard fly food (Genesee Scientific) and raised at 25°C. The *GMR*-GAL4 and *OK371*-GAL4 drivers were obtained from the Bloomington Stock Center. *MHC*-GAL4 was a gift from Guillermo Marques. All tested RNAi, deficiency, duplication, and classical allele lines are listed in supplemental table1.

#### Drosophila neuronal phenotype analysis

Synaptic bouton numbers from larvae (*n* = 10) were counted as previously described (*42*) with slight modification. Third-instar larvae for each group were collected and pinned on Sylgard dishes with tungsten pins. After dissecting the dorsal sides of larvae and removing the trachea and internal organs, the larvae were fixed with 4% paraformaldehyde in PBS. Larvae were stained with a presynaptic anti-HRP antibody (1:200, Jackson Immunoresearch) and postsynaptic anti-discs large antibody (1:200, Developmental Studies Hybridoma Bank). The NMJs at muscle 4 were used for all analyses.

#### Drosophila behavioral assay

Seven wandering third-instar larvae from each group were collected, washed, and placed onto a 3% agarose gel in a 10-cm dish. Larval crawling was recorded by a digital camera for 30 sec.

Moving distances of individual larva were tracked and measured by using Tracker software (Open Source Physics, 5.0.7). To test flight ability, 20 flies from each group were funneled into a 500-ml glass cylinder. The distribution of flies in the cylinder was recorded by a digital camera, and the average scores from five independent experiments were calculated.

#### Drosophila immunoblotting

Heads of adults were prepared and ground in LDS-sample buffer by using a motor-driven plastic pestle homogenizer and centrifuged at 16,000 *g* for 10 min. Supernatants were boiled for 10 min and analyzed by immunoblotting with the Odyssey FC system (LI-COR). Proteins were separated with 4%–20% ExpressPlus PAGE Gels (GenScript), transferred onto nitrocellulose membranes, and probed with anti-FLAG and anti-C2C10H antibodies. Anti-actin antibody was used as a loading control.

#### Adult muscle preparation and immunohistochemistry

The mitochondrial morphology and sarcomere structure of the indirect flight muscles were analyzed as previously described (*42*). Briefly, adult flies were fixed with 4% paraformaldehyde in PBS for 1 hour, embedded in OCT compound (Fisher Scientific), and frozen with liquid nitrogen. Samples were processed by a cryomicrotome (Leica). After fixing with 4% paraformaldehyde in PBS, samples were permeabilized with 0.2% Triton X-100 buffer in PBS and blocked with 5% BSA solution in PBS for 1 hour. Samples were incubated with streptavidin–Alexa Fluor 488 (Invitrogen) and phalloidin–Alexa Fluor 594 (Invitrogen) overnight at 4°C for mitochondrial and muscle staining, respectively. Samples were mounted with Prolong Diamond Antifade Reagent with DAPI and imaged with an LSM 710 confocal microscope (Zeiss) with 40× magnification.

#### ATP assay

Fly thoraxes from each group were collected and homogenized in 20 µl of homogenization buffer [100 mM Tris, 4 mM EDTA, and 6 M guanidine-HCL (pH 7.8)] and centrifuged at 16,000 *g* for 10 min. The supernatants were diluted 1:200 and 1:10 with deionized water and subjected to ATP concentration and protein concentration measurements, respectively. ATP concentration was determined by using the CellTiter Glo luminescent cell viability assay kit (Promega) and normalized to total protein.

#### Chemicals and peptides

CCCP was purchased from Sigma-Aldrich. The mitofusin agonists B1 and B/A-L were described previously (*49*). ScPM and PM1 peptides for TDP-43 were kindly provided by Xinglong Wang. Pink1_490 and Ub_60 peptides were synthesized by Thermo Scientific with *N*-terminal TAMRA 5/6 and *C*-terminal amidation.

#### DNA constructs

All cDNAs for human *CHCHD10^WT^* and variants were synthesized and inserted in the pcDNA3 vector containing a FLAG, HA, or Myc tag by Genescripts. *mTagRFP-T-Mito-7*, *EGFP-LC3*, *pEYFP-C1-DRP1*, *MFN2-YFP*, *pEYFP-N1-PINK1*, *LAMP1-mGFP*, *OTC*, and *ΔOTC* (i.e., amino acids 30–114 deleted) plasmids were obtained from Addgene. *FLAG-MFN2^WT^*, *FLAG-MFN2^S378A^*, and *FLAG-MFN2^S378D^* were described previously (*49*). *TARDBP^WT^*, *TARDBP^G298S^*, *TARDBP^A315T^*, and *TARDBP^A382T^* plasmids were gifts from Xinglong Wang.

#### CRISPR/Cas9-mediated gene editing and generation of cell lines

Two plasmid vectors [pSpCas9(BB)-2A-Puro (XP459) V2.0] containing an sgRNA oligomer for *CHCHD10* targeting near the *N*-terminal region were purchased from GenScript. Sequences for the sgRNA targeting the *N*-terminal of *CHCHD10* are as follows: 5′-GCCTCGGGGAAGCCGCAGCG and 5′-GCCGCGCTGCGGCTTCCCCG. The plasmids were transfected into HeLa and HeLa^YFP-Parkin^ cells with jetPrime reagent. After 24 hours, the transfected cells were detached with trypsin and plated individually into 96-well round-bottom plates. The cells were then expanded, and single clones were analyzed by immunofluorescence and immunoblotting to screen protein levels in *CHCHD10^KO^* cells.

#### Cell culture and transfection

Control HeLa cells, HeLa^YFP-Parkin^, HeLa with stable *PINK1* expression (HeLa^Pink1-V5/His^), and HeLa cells containing the V5/His backbone were gifts from Richard Youle. *PINK1^KO^* HeLa cells and those expressing the backbone construct were kindly provided by Wade Harper. Cells were maintained in culture in Dulbecco modified Eagle medium (Gibco) supplemented with 10% fetal bovine serum (Gibco), 1× penicillin/streptomycin (Invitrogen) and GlutaMax-1× (Gibco). Cells were transfected by using FuGENE 6 transfection reagent (Promega), Lipofectamine 3000 (Invitrogen), or jetPrime (Polyplus). RNAi-mediated knockdown of target genes was performed by transfection of the ON-TARGET plus-SMART pool siRNAs (Dharmacon) by using Lipofectamine RNAiMAX (Invitrogen) for the following genes: nontargeting control, *PINK1*, *MFN1*, *MFN2*, *OPTN*, *NDP52*, and *DRP1*. Pink1_490 and Ub_60 peptides were delivered by using PULSin protein delivery reagent (Polyplus).

#### Antibodies and immunoblotting

The following primary antibodies were used: FLAG (Sigma and Proteintech), FLAG-Alexa Fluor 488 (Invitrogen), HA (Cell Signaling and Proteintech), Myc (Proteintech), V5 (Life Technology), CHCHD10 (Proteintech), TOM20 (Cell Signaling and Santa Cruz Biotechnology), TDP-43 (Proteintech and Santa Cruz Biotechnology), HSP60 (Cell Signaling Technology), COX2 (Abcam), LC3B (Cell Signaling Technology), LAMP1 (Cell Signaling Technology), phospho-ubiquitin (EMD Millipore), Actin (Santa Cruz Biotechnology and Proteintech), PINK1 (Santa Cruz Biotechnology and Novus), DRP1 (Cell Signaling Technology), MFN1 (Cell Signaling Technology), MFN2 (Cell Signaling Technology), NDP52 (Proteintech), OPTN (Proteintech), CHOP (Cell Signaling Technology), and OTC (Novus). Rabbit anti-CG5010 (C2C10H) antibody was a gift from Nobutaka Hattori. Samples were collected and lysed in RIPA buffer (Cell Signaling Technology) containing a protease inhibitor cocktail (Sigma) and subsequently separated by SDS-PAGE after measuring protein concentration by BCA (Pierce). Immunoblots were visualized and analyzed with the Odyssey FC system (Li-Cor).

#### Immunofluorescence staining and imaging

Cells were plated on 4-well chamber slides (Lab Tek), fixed with 4% paraformaldehyde in PBS (EMS Millipore), permeabilized with 0.1% Triton X-100, and blocked with 5% bovine serum albumin (BSA) in PBS. Primary antibodies were diluted 1:100 to 1:250 in 5% BSA in PBS and incubated overnight at 4°C. Samples were then rinsed three times with PBS-tween and incubated with secondary antibodies for 1.5 hours at room temperature. Coverslips were mounted onto microscope slides with Prolong Diamond Antifade Reagent with DAPI (Invitrogen). Samples were observed with an LSM 710 confocal microscope (Zeiss). For live-cell imaging, HeLa cells were plated on glass-bottom dishes (Thermo Scientific) and imaged with an LSM 710 confocal microscope (Zeiss) equipped with an environmental chamber for controlling CO_2_ concentration and temperature (37°C).

#### Solubility and biochemical analyses

Transfected cells were washed twice with PBS, lysed in cold RIPA buffer [50 mM Tris (pH7.5), 150 mM NaCl, 1% Triton X-100, 0.5% sodium deoxycholate, 0.1% SDS, and 1 mM EDTA) and sonicated on ice. Cellular lysates were cleared by ultracentrifugation at 100,000 *g* for 30 min at 4°C to prepare RIPA-soluble fractions. RIPA-insoluble pellets were washed twice with protease inhibitors in cold PBS, sonicated, and recentrifuged. RIPA-insoluble proteins were extracted with urea buffer [7 M urea, 2 M thiourea, 4% CHAPS, 30 mM Tris (pH 8.5)], sonicated, and centrifuged at 100,000 *g* for 30 min at 22°C. Supernatants from the first centrifugation were analyzed for soluble fractions. Total protein in each sample was measured by the BCA method and resolved on ExpressPlus PAGE 4%–20 % gels (GenScript).

#### Cellular and mitochondrial fractionation

Mitochondria were isolated from cells with a mitochondrial isolation kit (Thermo Scientific) by using Dounce homogenizers. A mixture of cytosolic and mitochondrial fractions was obtained after low-speed centrifugation at 500 *g* for 15 min at 4°C. Mitochondrial-enriched pellets were collected at 3000 *g* for 10 min at 4°C. Cytosolic supernatants were obtained and cleared by centrifugation at 12,000 *g* for 10 min at 4°C. Cross-contamination between fractions was analyzed with anti-tubulin and anti-HSP60 antibodies for each cytosolic and mitochondrial compartment.

#### Mitochondrial respiratory activity assay

Mitochondrial respiration in HeLa, *CHCHD10^KO^* HeLa, HeLa^YFP-Parkin^, and *CHCHD10^KO^*/ HeLa^YFP-Parkin^ cells were measured by using the Seahorse Extracellular Flux Analyzer XFp (Agilent Technologies) with the XF Cell Mito Stress Test kit (Agilent Technologies). Transfected cells (1 × 10^4^) were plated into V3-PS 96-well plates the day before performing the assay. Assay media were supplemented with 1 mM pyruvate, 2 mM glutamine, and 15 mM glucose, with pH adjusted to 7.6. Standard mitochondria stress tests were performed by first measuring basal values followed by measurements after sequential addition of 1 µM oligomycin, 0.5 µM FCCP, and 0.5 µM rotenone/antimycin A. After the assay, protein concentrations of each well were determined via BCA assay and used to normalize oxygen consumption rate values.

---

**Fig. S1.**
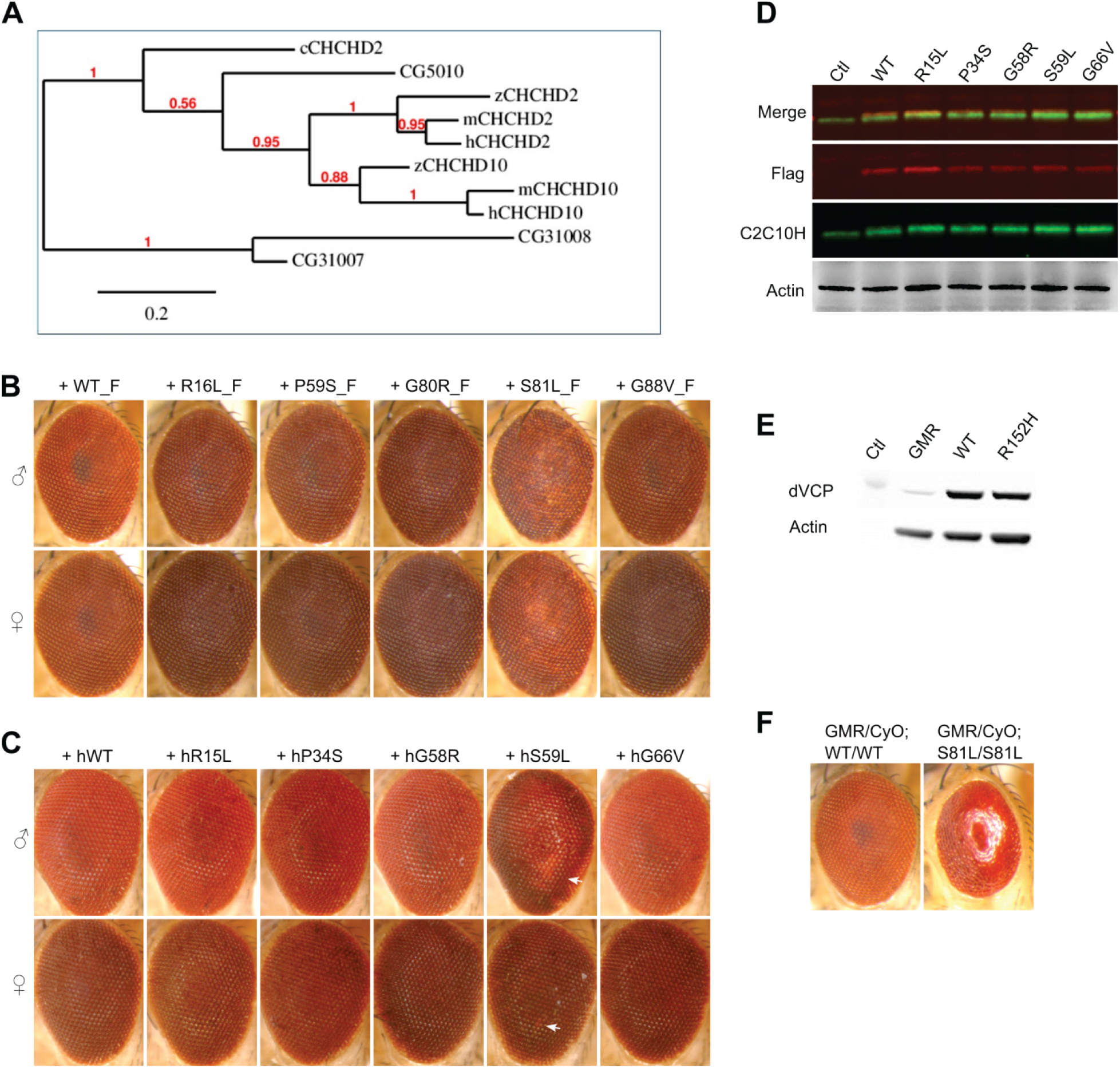
Eye phenotypes in *Drosophila* expressing *CHCHD2* and *CHCHD10.* (**A**) Phylogenetic tree for *CHCHD2* and *CHCHD10*. The genetic tree was generated with Phylogeny.fr. (**B**) FLAG-tagged *C2C10H^S81L^* causes age-dependent rough eye phenotypes in 40-day-old flies. (**C**) Expressing human *CHCHD10^S59L^* induces a mild rough eye phenotype in 40-day-old flies. Arrows indicate depigmented regions. (**D**) Proteins extracted from heads expressing FLAG-tagged *C2C10H* variants by *GMR*-GAL4 and subjected to immunoblotting for FLAG, C2C10H, and actin. (**E**) Exogenous expression of *TER94* with *GMR*-GAL4. (**F**) Expression from two copies of *C2C10H^S81L^* causes a severe rough eye phenotype at eclosion.

**Fig. S2.**
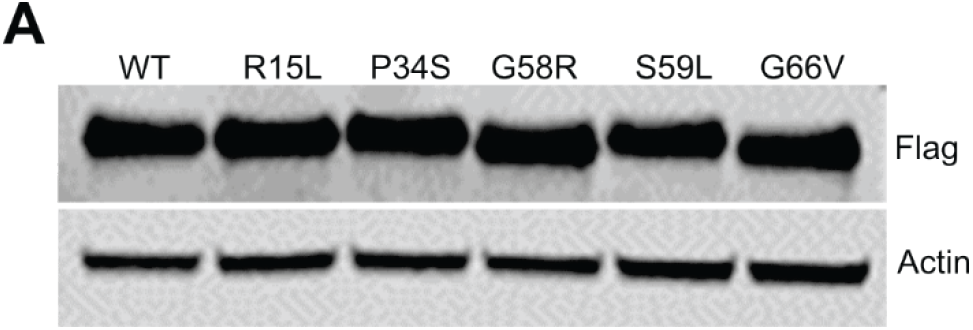
Expression of WT and mutant *CHCHD10* in HeLa cells. (**A**) Expression of FLAG-tagged *CHCHD10^WT^* and variants in HeLa cells. HeLa cells were transiently transfected and subjected to immunoblotting with anti-FLAG and anti-actin antibodies.

**Fig. S3.**
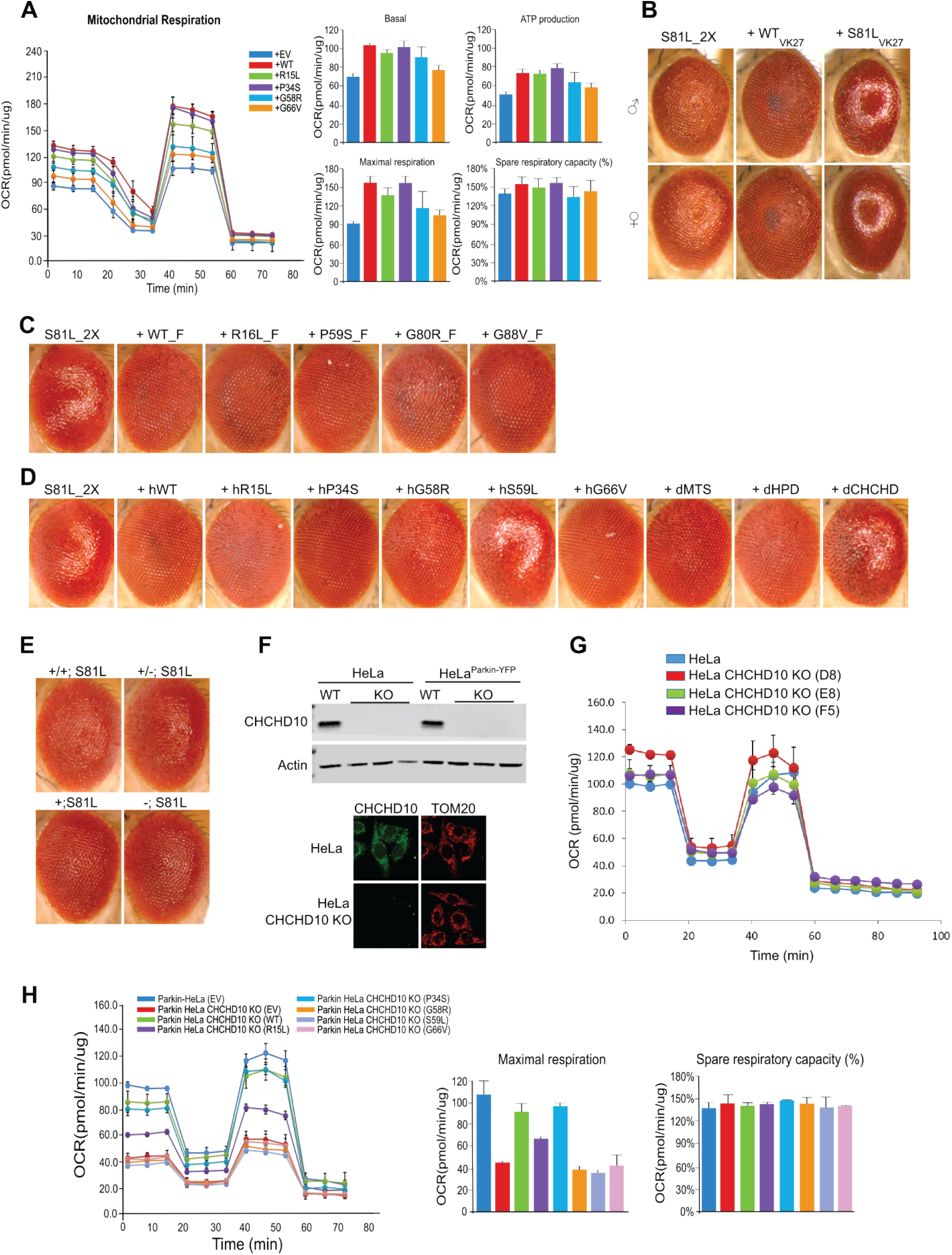
Co-expression of WT and mutant *CHCHD10* constructs in *Drosophila* and HeLa cells. (**A**) HeLa cells were co-transfected with FLAG-tagged *CHCHD10^S59L^* and FLAG-tagged *CHCHD10^WT^* or variants. Empty vector (EV) was used as a control. After 24 hours, mitochondrial respiration for each group was measured by Seahorse XF Cell Mito Stress tests. Data shown are mean ± SD (*n* = 3 technical replicates). (**B**) Expression of *C2C10H^WT^* inserted in the *VK27* site improved *C2C10H^S81L^*-induced rough eye phenotypes, whereas insertion of *C2C10H^S81L^* in the *VK27* site exacerbated the phenotypes. (**C**) Expression of mutant *C2C10H* inserted in the *VK27* site improved *C2C10H^S81L^*-induced rough eye phenotypes. (**D**) Expression of human *CHCHD10^WT^* and variants mitigated *C2C10H^S81L^*-induced rough eye phenotypes. (**E**) *C2C10H^S81L^*-induced rough eye phenotypes were robust in the *C2C10H^null^* background. (**F**) Lysates from HeLa cells, three *CHCHD10^KO^* HeLa cell lines (C10 KO), HeLa^YFP-Parkin^ cells, and three *CHCHD10^KO^*/HeLa^YFP-Parkin^ cell lines were immunoblotted with an anti-CHCHD10 antibody (upper panel) and immunostained with anti-CHCHD10 and anti-TOM20 antibodies (lower panel). (**G**) Mitochondrial respiration of HeLa cells and three *CHCHD10^KO^* HeLa cell lines was measured by Seahorse XF Cell Mito Stress tests. Data shown are mean ± SD (*n* = 3 technical replicates). (**H**) *CHCHD10^KO^*/HeLa^YFP-Parkin^ cells were transfected with FLAG-tagged *CHCHD10^WT^* and variants. After 24 hours, mitochondrial respiration was measured by Seahorse XF Cell Mito Stress tests.

**Fig. S4.**
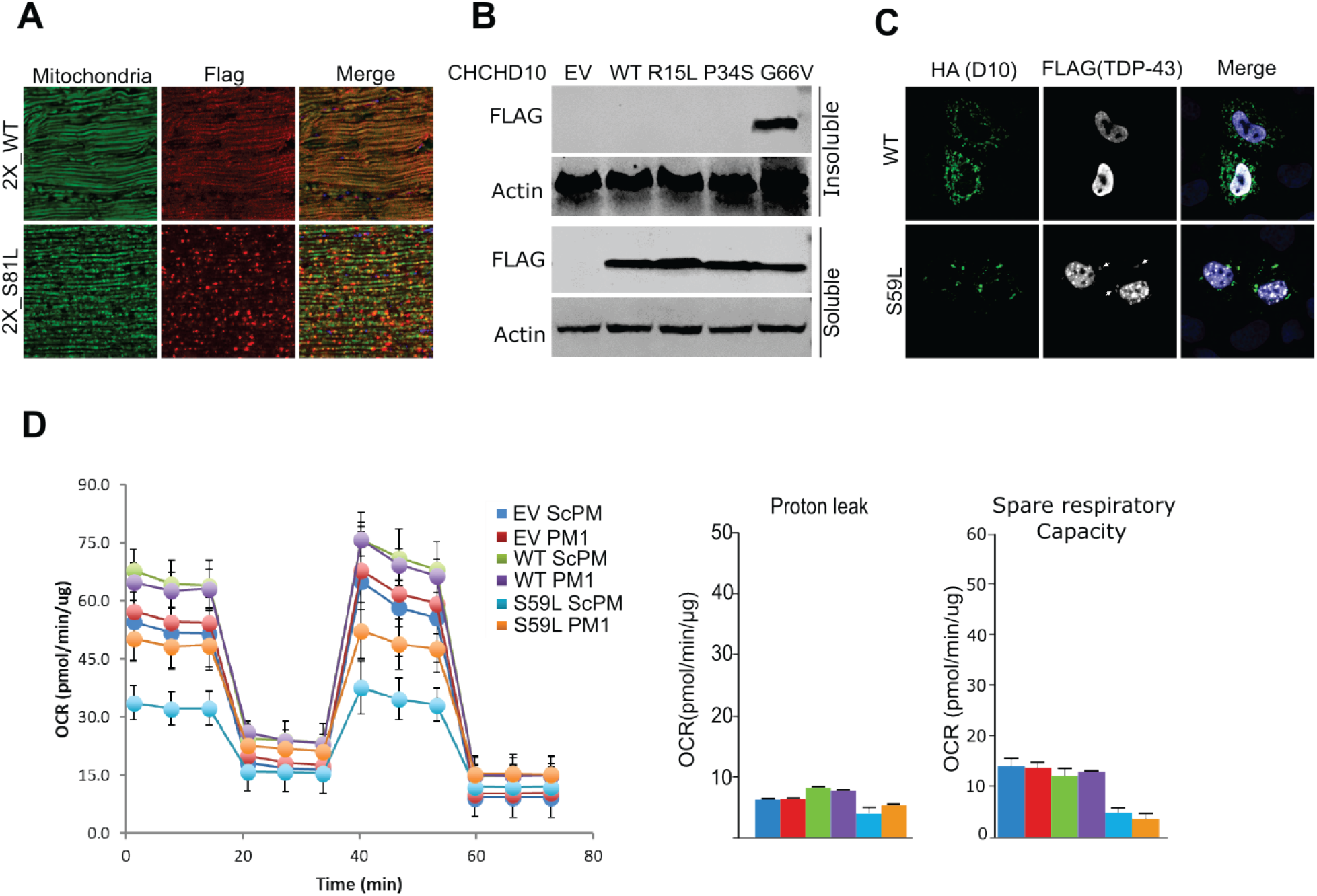
Mitochondrial localization and activity in WT and mutant *CHCHD10* flies and HeLa cells. (**A**) Indirect flight muscles from 10-day-old *MHC*-GAL4>*UAS*-2X *C2C10H^WT^* and 2X_*C2C10H^S81L^* males were immunostained with streptavidin–Alexa Fluor 488 (green) and anti-FLAG antibody (red) to visualize mitochondria and C2C10H^WT^ or C2C10H^S81L^, respectively. (**B**) HeLa cells were transfected with FLAG-tagged *CHCHD10^WT^* and variants. After 24 hours, cells were subjected to sequential protein extraction with RIPA and urea buffers. Immunoblotting was conducted with anti-FLAG or anti-actin (loading control) antibodies. (**C**) HeLa cells were co-transfected with FLAG-tagged *TARDBP* and HA-tagged *CHCHD10^WT^* or *CHCHD10^S59L^*. After 24 hours, cells were stained with anti-FLAG and anti-HA antibodies. Arrows indicate mitochondrial localization of TDP-43. (**D**) Mitochondrial respiration was measured by Seahorse XF Cell Mito Stress tests.

**Fig. S5.**
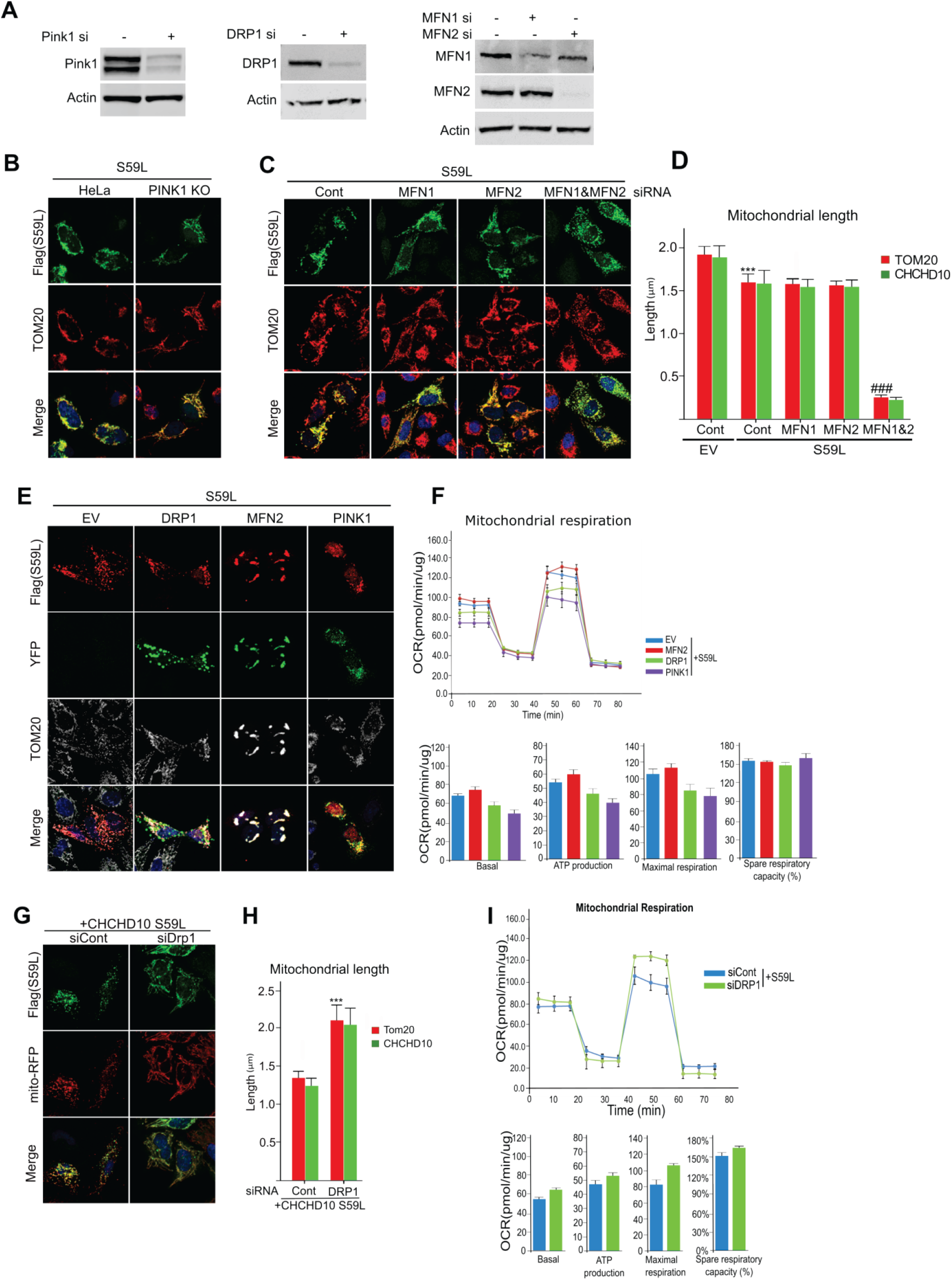
Role of the PINK1/parkin pathway on the *CHCHD10^S59L^*-induced phenotype in HeLa cells. (**A**) HeLa cells were transfected with siRNAs targeting *PINK1*, *DRP1*, *MFN1*, or *MFN2* or control siRNA. Immunoblotting confirmed successful knockdown of target genes. (**B**) HeLa cells and *PINK1^KO^* HeLa cells were transfected with FLAG-tagged *CHCHD10^S59L^* or empty vector (EV). Representative images of transfected HeLa cells were immunostained with antibodies against FLAG (green, CHCHD10) and TOM20 (red, mitochondria). (**C**) HeLa cells were transfected with siRNAs targeting *MFN1* and/or *MFN2*. The cells were transfected with FLAG-tagged *CHCHD10^S59L^* 24 hours after siRNA transfection. Representative images of transfected HeLa cells immunostained with antibodies against FLAG (green, CHCHD10) and TOM20 (red, mitochondria). (**D**) Quantification of TOM20 and CHCHD10^S59L^-FLAG signal strength. Data shown are mean ± SD (one-way ANOVA followed by posthoc Tukey analysis, \*\*\**P* < 0.001 *vs.* EV; *n* = 3 biological replicates; ###*P* < 0.001 *vs. CHCHD10^S59L^*-only cells, with at least 15 cells in each group). (**E**) HeLa cells were co-transfected with YFP-tagged *DRP1*, *MFN2*, or *PINK1* and FLAG-tagged *CHCHD10^S59L^*. Transfected HeLa cells were immunostained with an antibody against FLAG (red, CHCHD10) and TOM20 (gray, mitochondria). (**F**) Mitochondrial respiration was measured by Seahorse XF Cell Mito Stress tests 24 hours after transfection. (**G**) HeLa cells were transfected with a siRNA targeting *DRP1*. After 18 hours of siRNA transfection, the cells were transfected with FLAG-tagged *CHCHD10^S59L^*. Representative images of transfected HeLa cells immunostained with antibodies against FLAG (red, CHCHD10) and TOM20 (gray, mitochondria). (**H**) Quantification of TOM20 and CHCHD10^S59L^-FLAG signal strength. Data shown are mean ± SD (one-way ANOVA followed by posthoc Tukey analysis, \*\*\**P* < 0.0001 *vs.* EV; *n* = 3 biological replicates). (**I**) Mitochondrial respiration was measured by Seahorse XF Cell Mito Stress tests at 24 hours after transfection. Data shown are mean ± SD (*n* = 3 technical replicates).

**Fig. S6.**
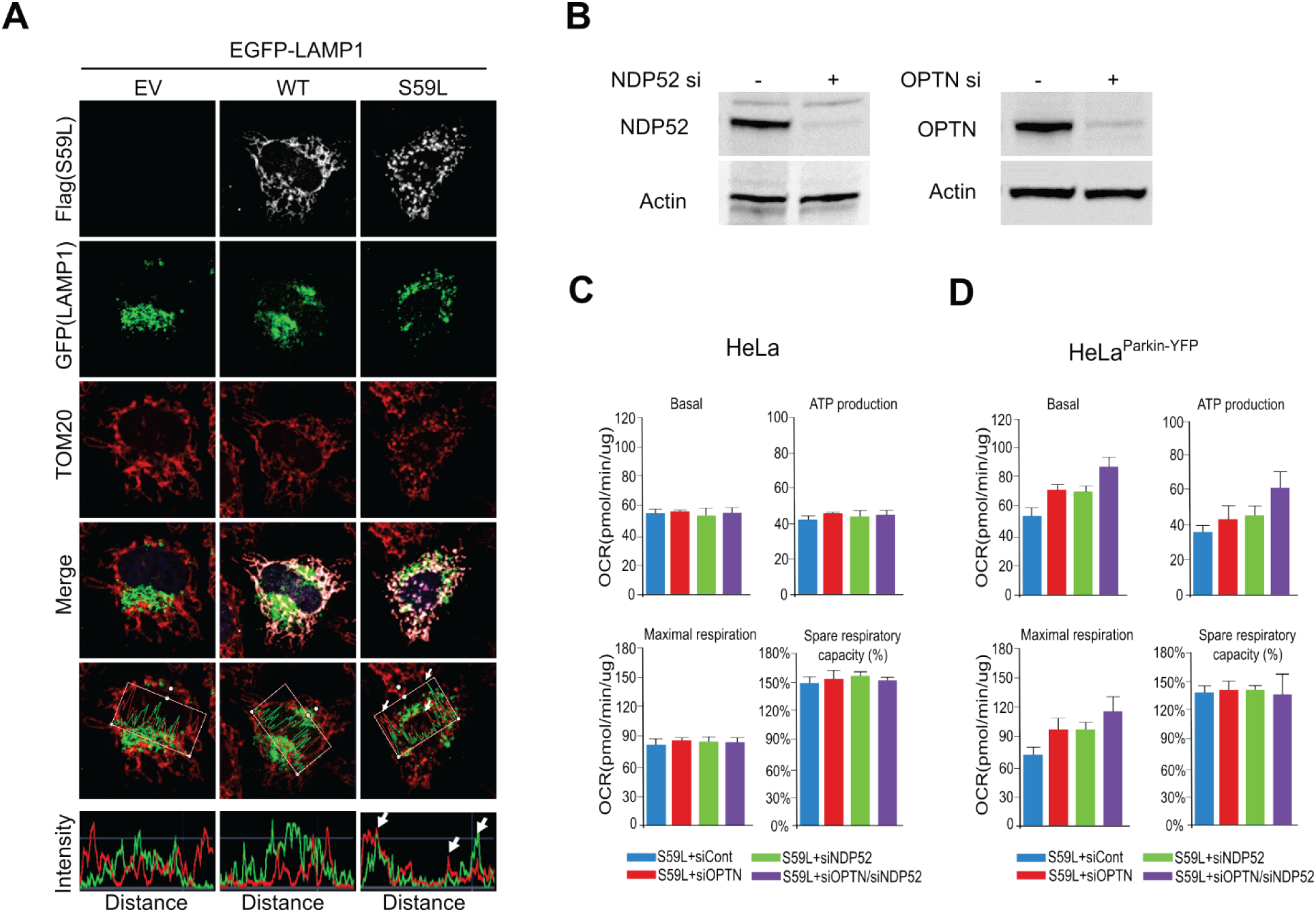
Effect of *CHCHD10^S59L^* on mitophagy. (**A**) HeLa cells were transfected with *EGFP-LAMP1* and FLAG-tagged *CHCHD10^S59L^*. After 24 hours of transfection, the cells were immunostained with antibodies against FLAG and TOM20. Graphs show the fluorescence intensity profiles of EGFP-LAMP1 and TOM20 along the regions boxed in red. Arrows indicate highly merged LAMP1–CHCHD10^S59L^ region. (**B**) HeLa cells were transfected with siRNAs targeting *NDP52* or *OPTN* or control siRNA. Immunoblotting confirmed RNAi-mediated knockdown of target genes. (**C**) HeLa cells and (**D**) HeLa^YFP-Parkin^ cells were transfected with siRNAs targeting *NDP52* and/or and *OPTN*. Mitochondrial respiration was measured by Seahorse XF Cell Mito Stress tests at 24 hours after transfection.

**Fig. S7.**
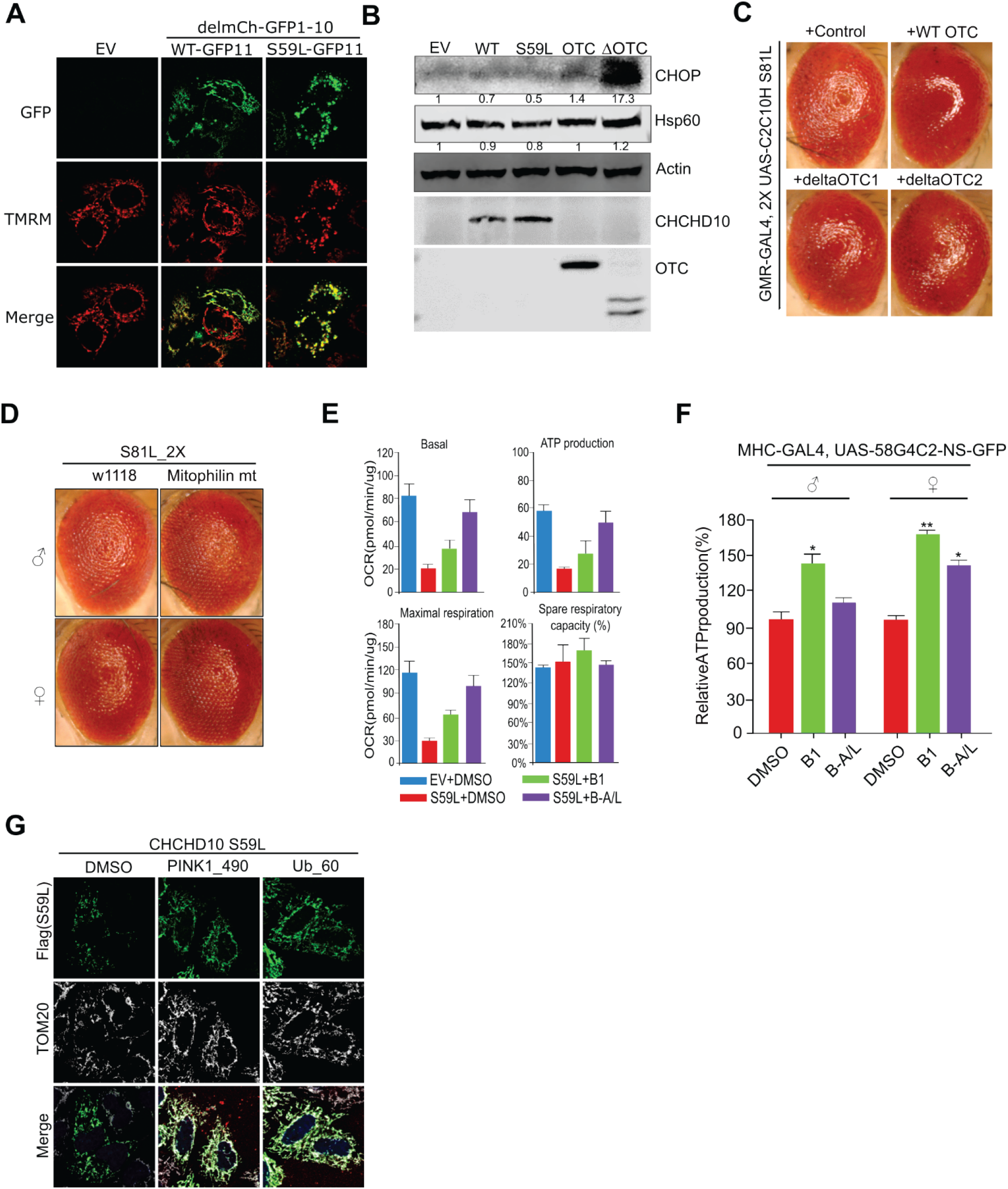
Effect of *CHCHD10^S59L^* on mitochondrial membrane potential and the UPR. (**A**) HeLa cells were transfected with GFP11-tagged *CHCHD10^WT^* or *CHCHD10^S59L^* and mitochondrial-localized GFP1–10. After 24 hours, cells were incubated with TMRM (100 nM). (**B**) HeLa cells transfected with *CHCHD10^WT^*, *CHCHD10^S59L^*, or *OTC* were analyzed by immunoblotting for CHOP and HSP60 proteins. (**C**) Expression of *OTC* or *ΔOTC* did not affect the rough eye phenotype induced by *C2C10H^S81L^*. (**D**) Overexpression of mutant *mitofilin* partially rescued *C2C10H^S81L^*-induced rough eye phenotypes. (**E**) HeLa cells were transfected with FLAG-tagged *CHCHD10^S59L^* and treated with the MFN2 agonists B1 (50 nM) or B-A/L (5 nM) for 24 hours. Mitochondrial respiration was measured with the Seahorse XF Cell Mito Stress test kit. (**F**) ATP levels in thoraxes from 10-day-old flies expressing *C9orf72* with expanded GGGGCC repeats fed with MFN2 agonists (each 10 μM). DMSO (0.1%) was used as a vehicle control. (**G**) HeLa cells were pre-transfected for 24 hours with FLAG-tagged *CHCHD10^S59L^* and then treated with the peptide inhibitors PINK1_490 (0.15 µg/ml) and Ub_60 (0.15 µg/ml). DMSO was used as a vehicle control. HeLa cells were visualized with antibodies against FLAG and TOM20.

**Fig. S8.**
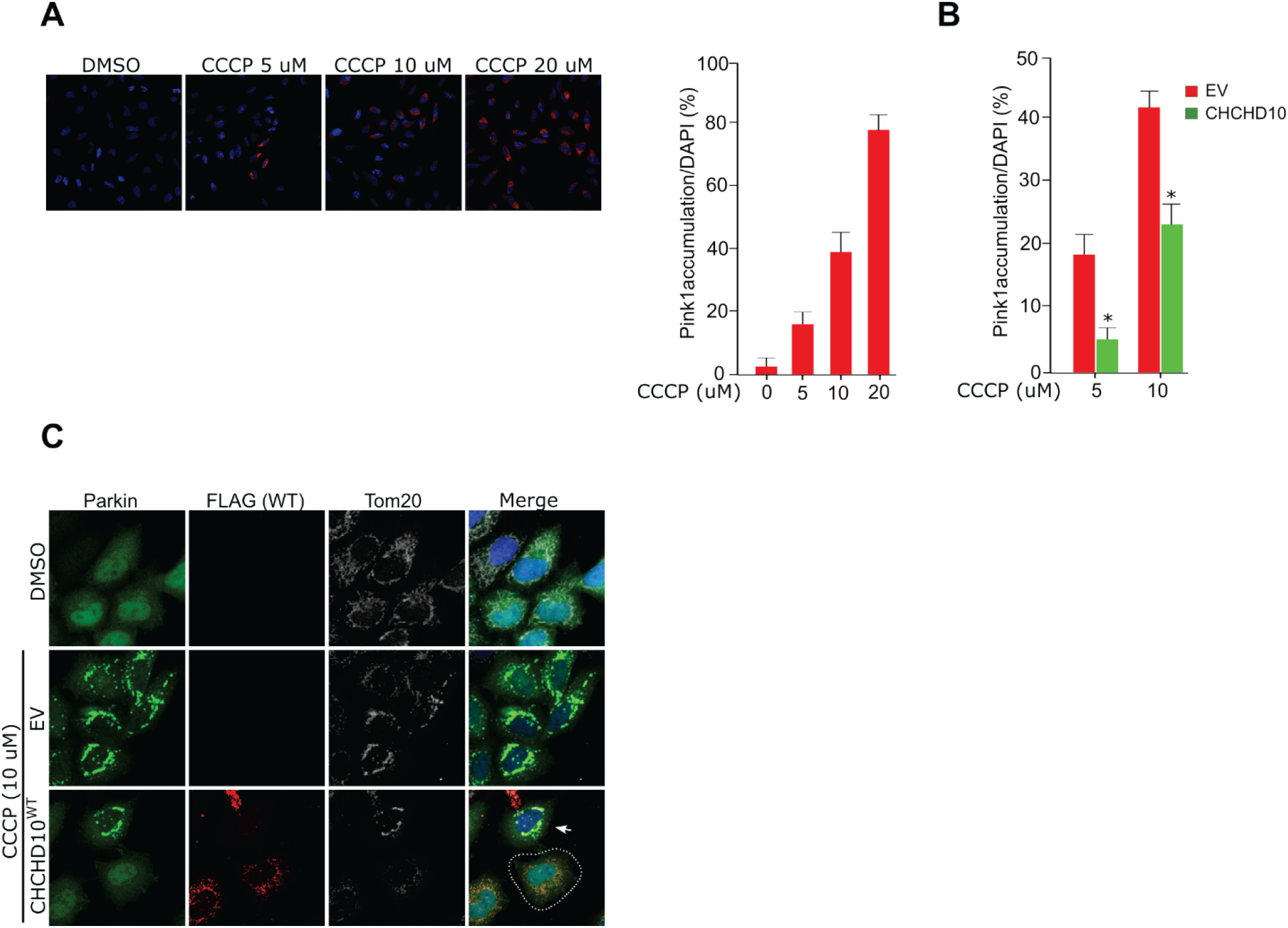
Effect of *CHCHD10* on CCCP-induced PINK1/parkin accumulation. (**A**) HeLa^PINK1-V5-His^ cells treated with multiple concentrations of CCCP for 6 hours were analyzed with a V5 antibody (red) and DAPI (blue). The percentage of PINK1-positive cells was calculated from the number of DAPI^+^ cells. For each sample, at least 300 cells were counted (*n* = 3 biological replicates). (**B**) The percentage of PINK1-positive cells from empty vector (EV)-or *CHCHD10^WT^*-transfected cells were calculated after 5 or 10 μM CCCP treatment for 6 hours. Data are mean ± SD (one-way ANOVA, \**P* < 0.05; *n* = 3 independent experiments). (**C**) HeLa^YFP-Parkin^ cells transfected with EV or FLAG-tagged *CHCHD10^WT^* were treated with CCCP (10 μM) for 6 hours. Cells were analyzed with anti-FLAG (red) antibody, YFP (green), and DAPI (blue, nucleus) to visualize CHCHD10 and parkin proteins. Arrow indicates parkin accumulated in nontransfected cell neighboring a *CHCHD10*-transfected cell (white dashed line)

**Table S1.**
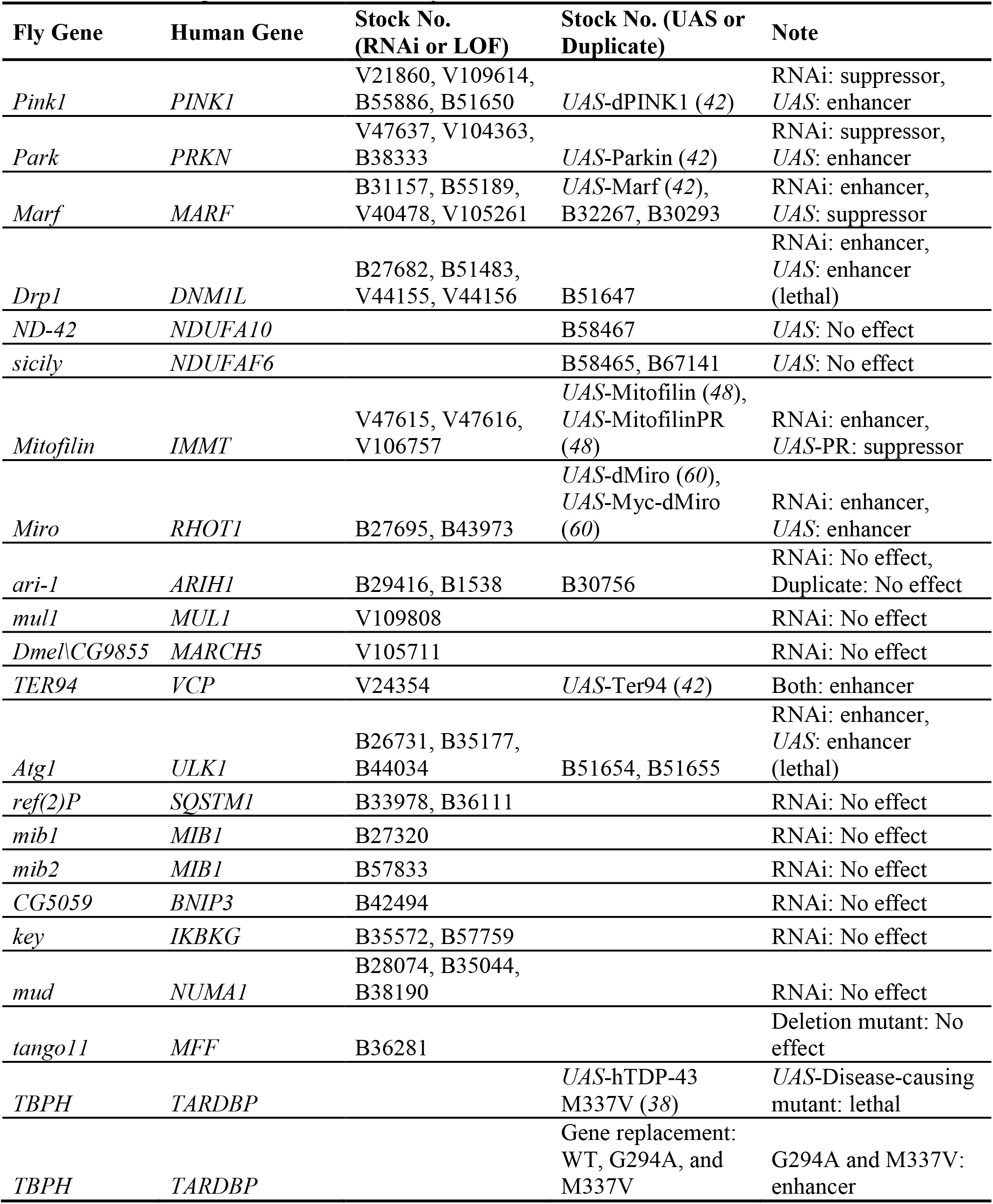
Drosophila lines used in study.

