## Supplemental information for "Dominant toxicity of ALS–FTD-associated *CHCHD10^S59L^* is mediated by TDP-43 and PINK1"

**Figure S1**

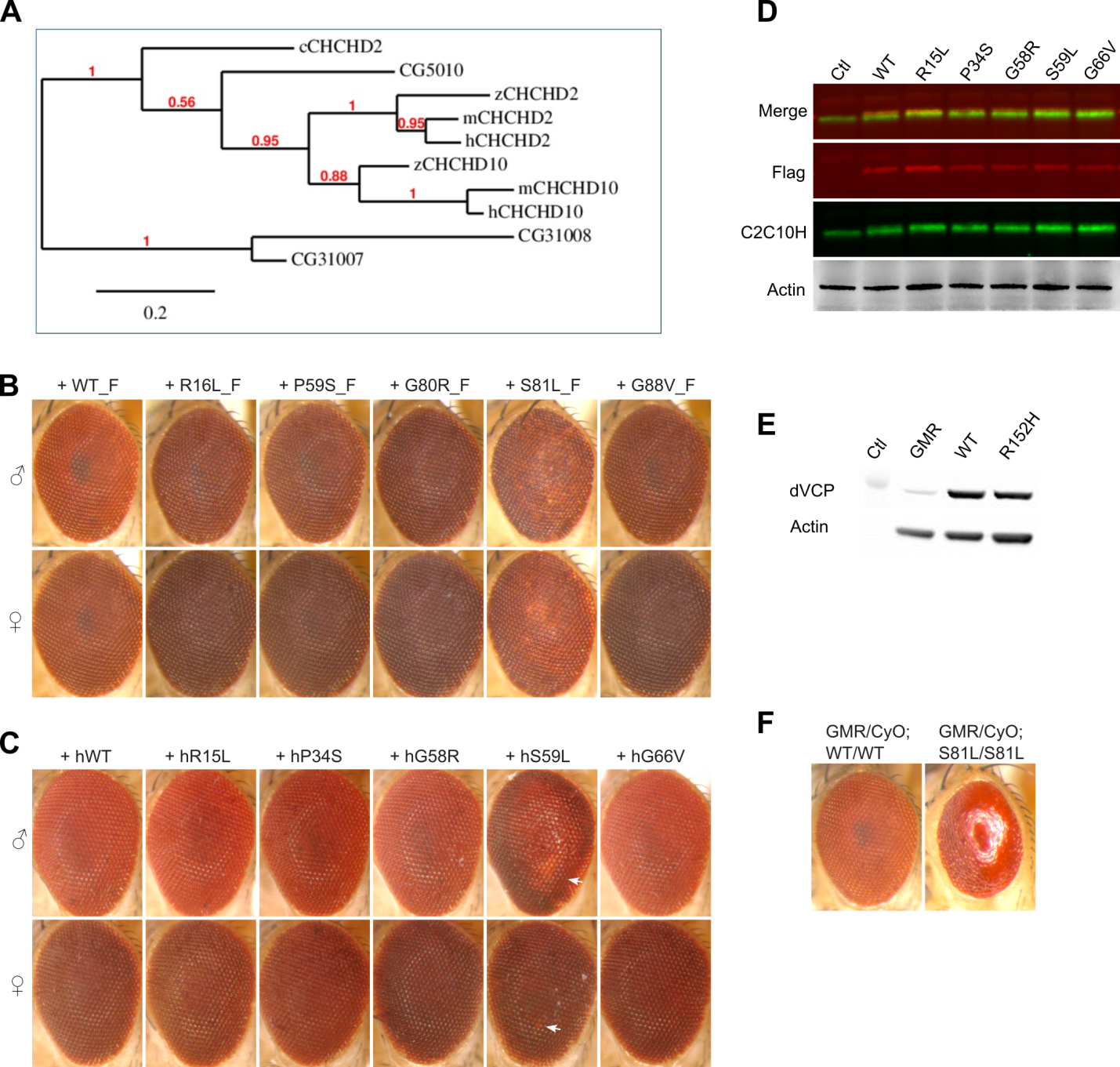

**Fig. S1. Eye phenotypes in *Drosophila* expressing *CHCHD2* and *CHCHD10.*** (**A**) Phylogenetic tree for *CHCHD2* and *CHCHD10*. The genetic tree was generated with Phylogeny.fr. (**B**) FLAG-tagged *C2C10H^S81L^* causes age-dependent rough eye phenotypes in 40-day-old flies. (**C**) Expressing human *CHCHD10^S59L^* induces a mild rough eye phenotype in 40-day-old flies. Arrows indicate depigmented regions. (**D**) Proteins extracted from heads expressing FLAG-tagged *C2C10H* variants by *GMR*-GAL4 and subjected to immunoblotting for FLAG, C2C10H, and actin. (**E**) Exogenous expression of *TER94* with *GMR*-GAL4. (**F**) Expression from two copies of *C2C10H^S81L^* causes a severe rough eye phenotype at eclosion.

**Figure S2**

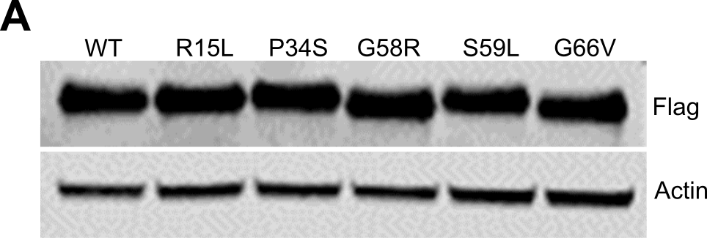

**Fig. S2. Expression of WT and mutant *CHCHD10* in HeLa cells.** (**A**) Expression of FLAG-tagged *CHCHD10^WT^* and variants in HeLa cells. HeLa cells were transiently transfected and subjected to immunoblotting with anti-FLAG and anti-actin antibodies.

**Figure S3**

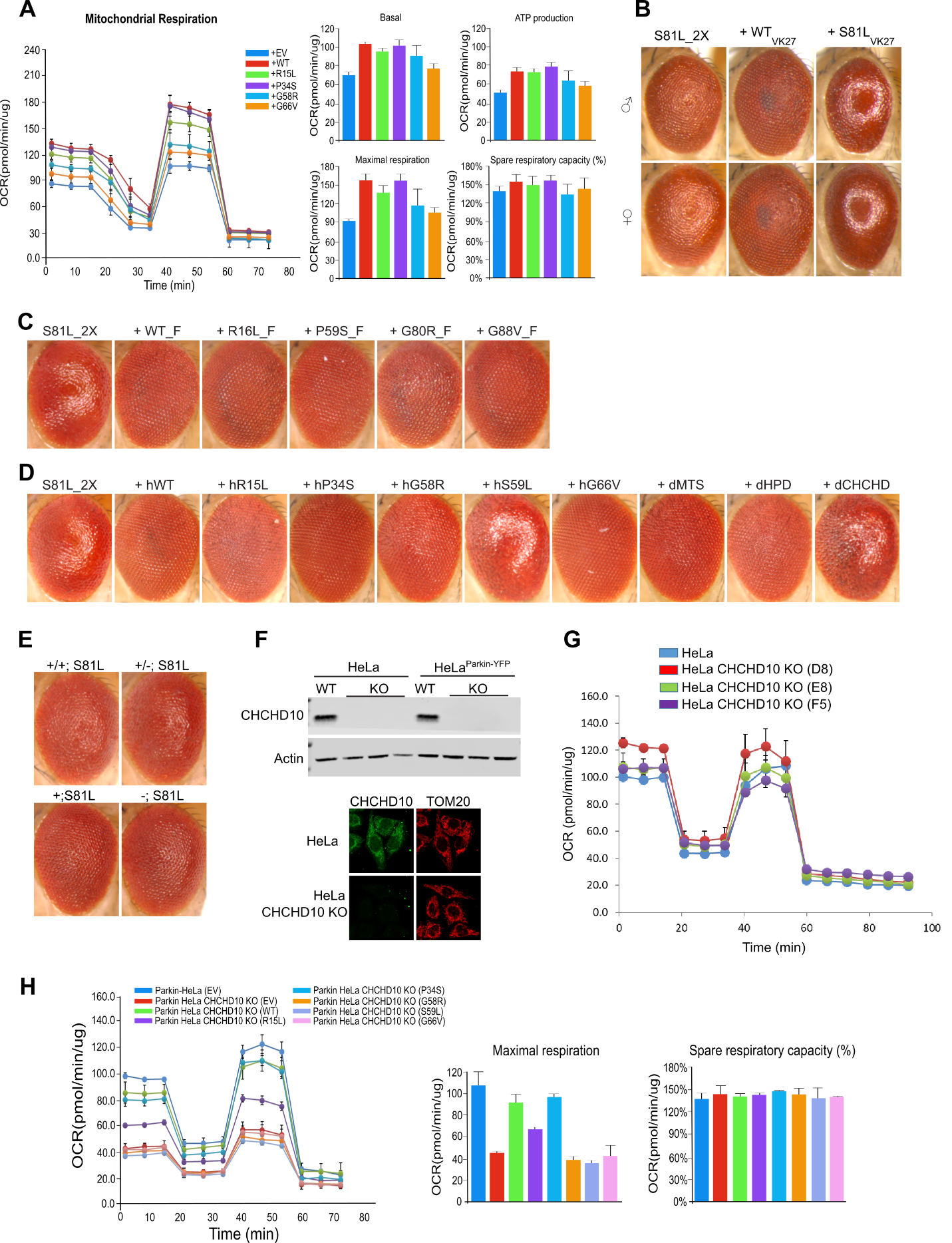

**Fig. S3. Co-expression of WT and mutant *CHCHD10* constructs in *Drosophila* and HeLa cells.** (**A**) HeLa cells were co-transfected with FLAG-tagged *CHCHD10^S59L^* and FLAG-tagged *CHCHD10^WT^* or variants. Empty vector (EV) was used as a control. After 24 hours, mitochondrial respiration for each group was measured by Seahorse XF Cell Mito Stress tests. Data shown are mean ± SD (*n* = 3 technical replicates). (**B**) Expression of *C2C10H^WT^* inserted in the *VK27* site improved *C2C10H^S81L^*-induced rough eye phenotypes, whereas insertion of *C2C10H^S81L^* in the *VK27* site exacerbated the phenotypes. (**C**) Expression of mutant *C2C10H* inserted in the *VK27* site improved *C2C10H^S81L^*-induced rough eye phenotypes. (**D**) Expression of human *CHCHD10^WT^* and variants mitigated *C2C10H^S81L^*-induced rough eye phenotypes. (**E**) *C2C10H^S81L^*-induced rough eye phenotypes were robust in the *C2C10H^null^* background. (**F**) Lysates from HeLa cells, three *CHCHD10^KO^* HeLa cell lines (C10 KO), HeLa^YFP-Parkin^ cells, and three *CHCHD10^KO^*/HeLa^YFP-Parkin^ cell lines were immunoblotted with an anti-CHCHD10 antibody (upper panel) and immunostained with anti-CHCHD10 and anti-TOM20 antibodies (lower panel). (**G**) Mitochondrial respiration of HeLa cells and three *CHCHD10^KO^* HeLa cell lines was measured by Seahorse XF Cell Mito Stress tests. Data shown are mean ± SD (*n* = 3 technical replicates). (**H**) *CHCHD10^KO^*/HeLa^YFP-Parkin^ cells were transfected with FLAG-tagged *CHCHD10^WT^* and variants. After 24 hours, mitochondrial respiration was measured by Seahorse XF Cell Mito Stress tests.

**Figure S4**

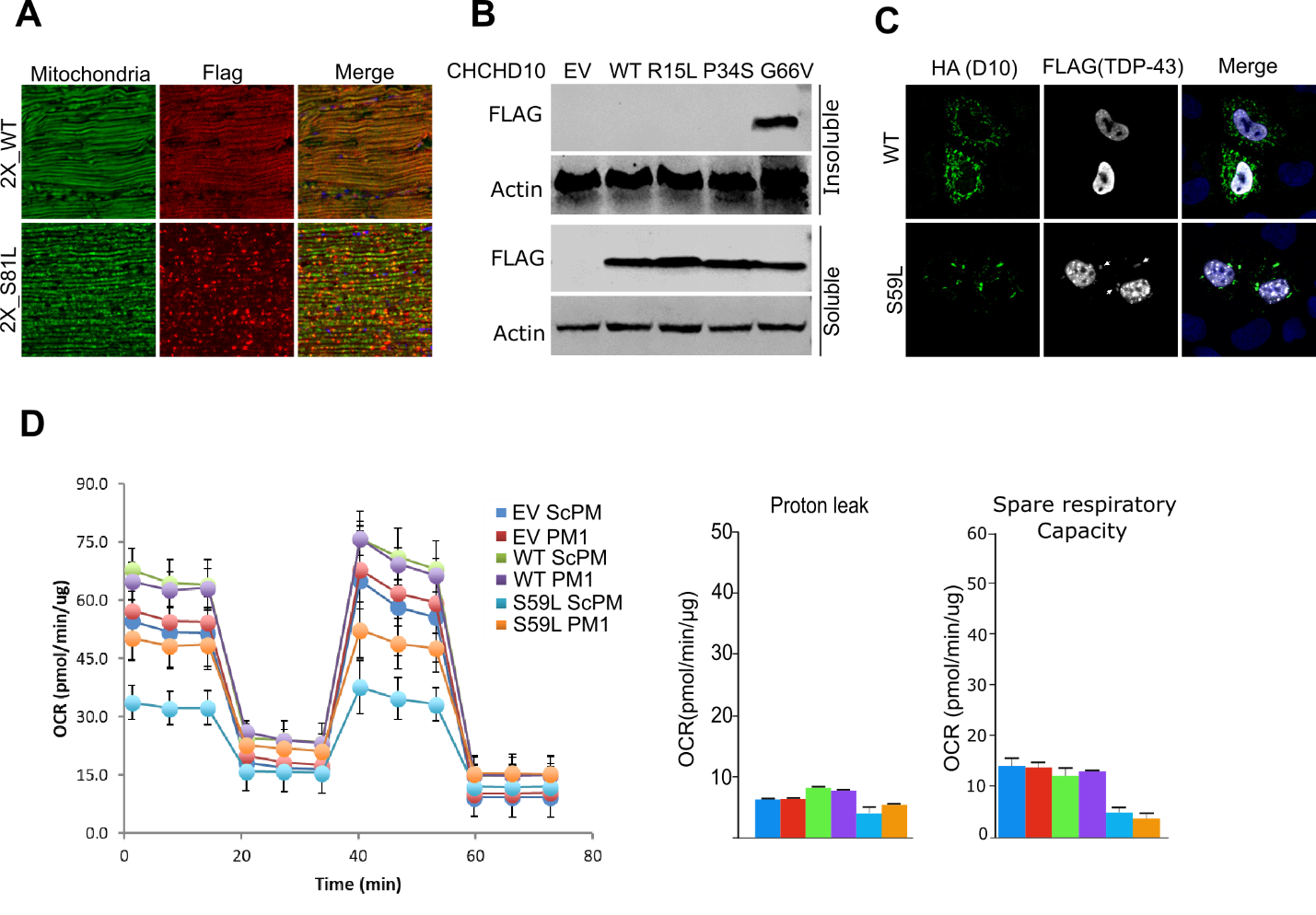

**Fig. S4. Mitochondrial localization and activity in WT and mutant *CHCHD10* flies and HeLa cells.** (**A**) Indirect flight muscles from 10-day-old *MHC*-GAL4>*UAS*-2X *C2C10H^WT^* and 2X_*C2C10H^S81L^* males were immunostained with streptavidin–Alexa Fluor 488 (green) and anti-FLAG antibody (red) to visualize mitochondria and C2C10H^WT^ or C2C10H^S81L^, respectively. (**B**) HeLa cells were transfected with FLAG-tagged *CHCHD10^WT^* and variants. After 24 hours, cells were subjected to sequential protein extraction with RIPA and urea buffers. Immunoblotting was conducted with anti-FLAG or anti-actin (loading control) antibodies. (**C**) HeLa cells were co-transfected with FLAG-tagged *TARDBP* and HA-tagged *CHCHD10^WT^* or *CHCHD10^S59L^*. After 24 hours, cells were stained with anti-FLAG and anti-HA antibodies. Arrows indicate mitochondrial localization of TDP-43. (**D**) Mitochondrial respiration was measured by Seahorse XF Cell Mito Stress tests.

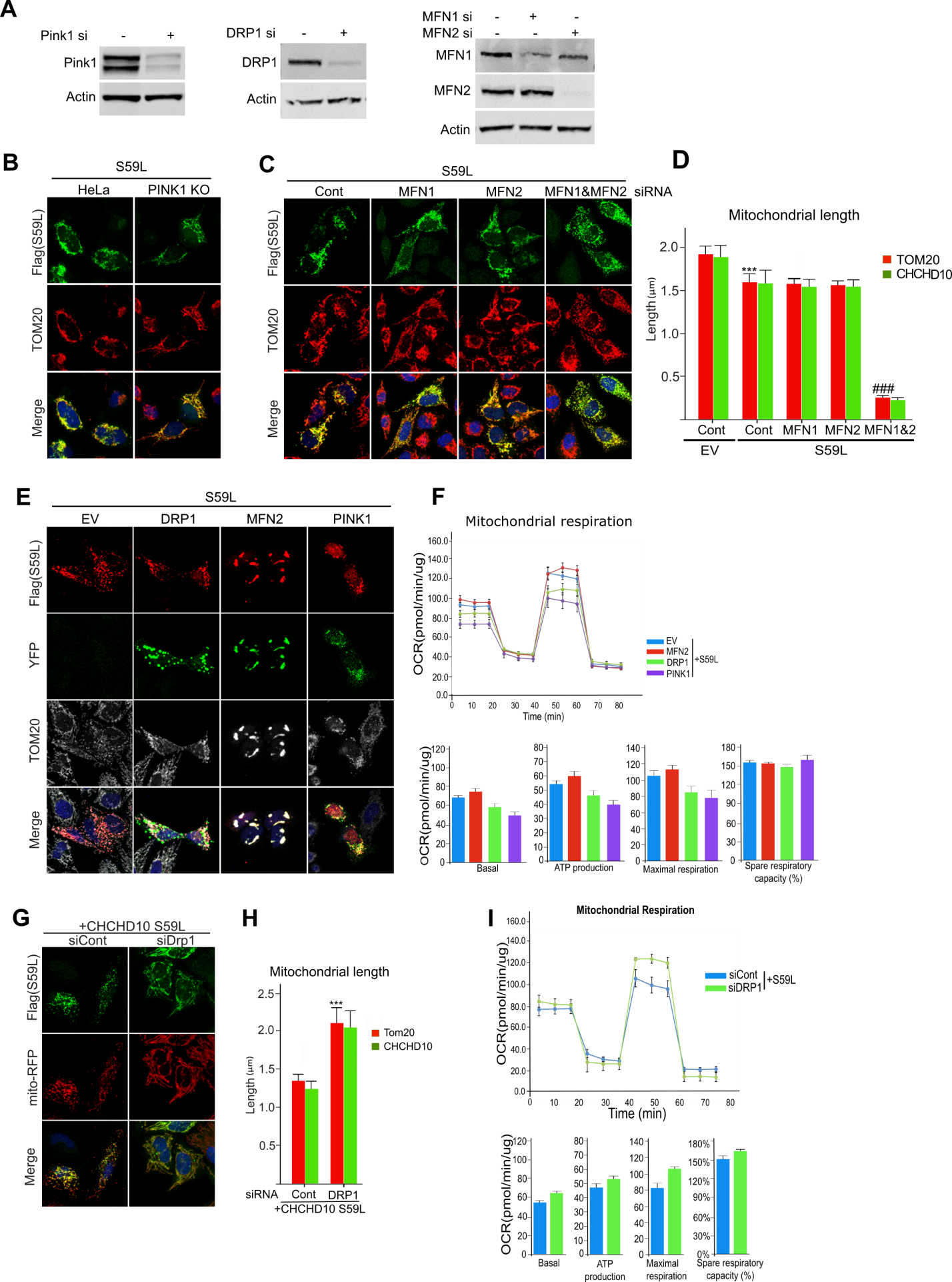
**Figure S5**

**Fig. S5. Role of the PINK1/parkin pathway on the *CHCHD10^S59L^*-induced phenotype in HeLa cells.** (**A**) HeLa cells were transfected with siRNAs targeting *PINK1*, *DRP1*, *MFN1*, or *MFN2* or control siRNA. Immunoblotting confirmed successful knockdown of target genes. (**B**) HeLa cells and *PINK1^KO^* HeLa cells were transfected with FLAG-tagged *CHCHD10^S59L^* or empty vector (EV). Representative images of transfected HeLa cells were immunostained with antibodies against FLAG (green, CHCHD10) and TOM20 (red, mitochondria). (**C**) HeLa cells were transfected with siRNAs targeting *MFN1* and/or *MFN2*. The cells were transfected with FLAG-tagged *CHCHD10^S59L^* 24 hours after siRNA transfection. Representative images of transfected HeLa cells immunostained with antibodies against FLAG (green, CHCHD10) and TOM20 (red, mitochondria). (**D**) Quantification of TOM20 and CHCHD10^S59L^-FLAG signal strength. Data shown are mean ± SD (one-way ANOVA followed by posthoc Tukey analysis, ****P* < 0.001 *vs.* EV; *n* = 3 biological replicates; ###*P* < 0.001 *vs.* *CHCHD10^S59L^*-only cells, with at least 15 cells in each group). (**E**) HeLa cells were co-transfected with YFP-tagged *DRP1*, *MFN2*, or *PINK1* and FLAG-tagged *CHCHD10^S59L^*. Transfected HeLa cells were immunostained with an antibody against FLAG (red, CHCHD10) and TOM20 (gray, mitochondria). (**F**) Mitochondrial respiration was measured by Seahorse XF Cell Mito Stress tests 24 hours after transfection. (**G**) HeLa cells were transfected with a siRNA targeting *DRP1*. After 18 hours of siRNA transfection, the cells were transfected with FLAG-tagged *CHCHD10^S59L^*. Representative images of transfected HeLa cells immunostained with antibodies against FLAG (red, CHCHD10) and TOM20 (gray, mitochondria). (**H**) Quantification of TOM20 and CHCHD10^S59L^-FLAG signal strength. Data shown are mean ± SD (one-way ANOVA followed by posthoc Tukey analysis, ****P* < 0.0001 *vs.* EV; *n* = 3 biological replicates). (**I**) Mitochondrial respiration was measured by Seahorse XF Cell Mito Stress tests at 24 hours after transfection. Data shown are mean ± SD (*n* = 3 technical replicates).

**Figure S6**

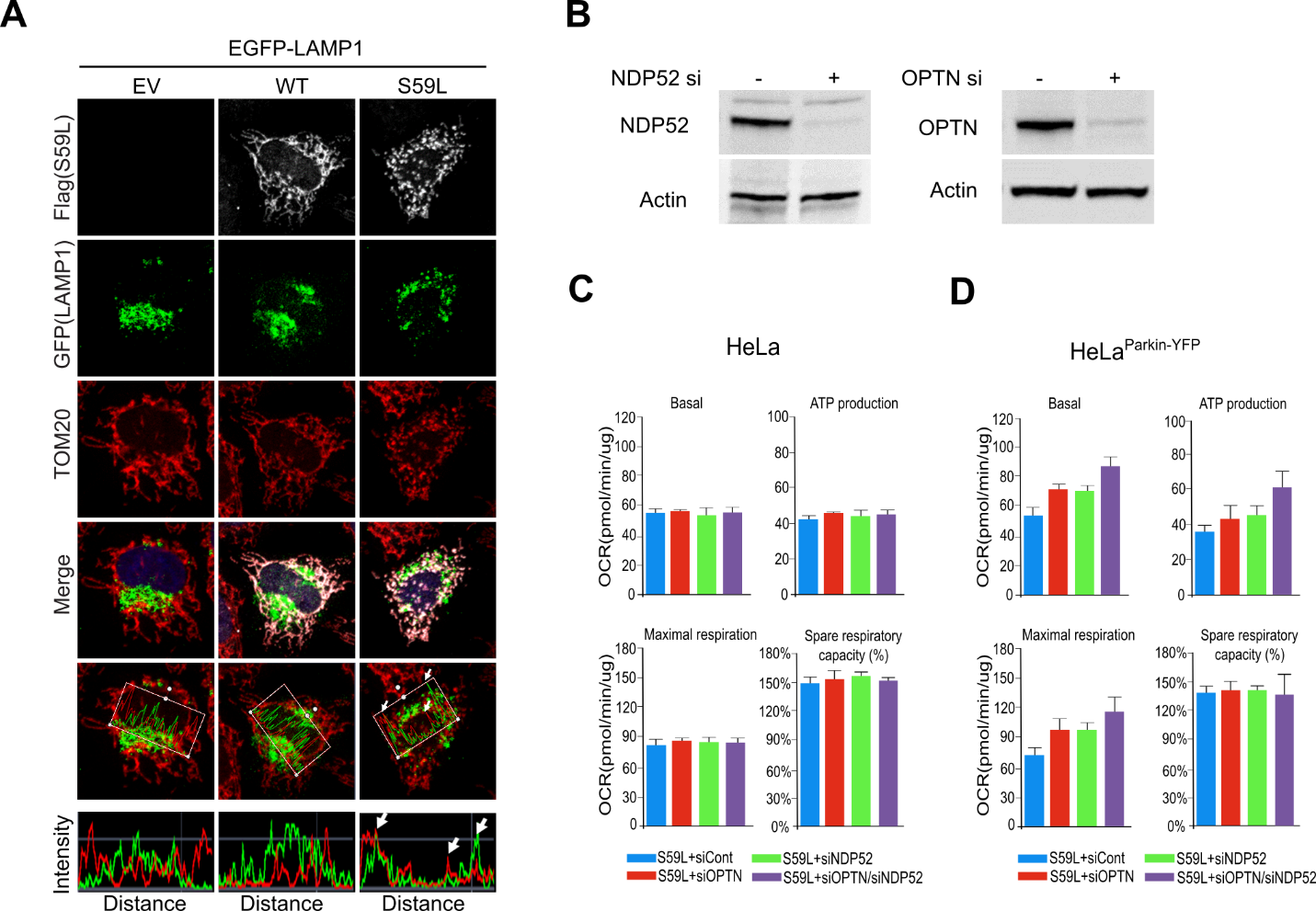

**Fig. S6. Effect of *CHCHD10^S59L^* on mitophagy.** (**A**) HeLa cells were transfected with *EGFP-LAMP1* and FLAG-tagged *CHCHD10^S59L^*. After 24 hours of transfection, the cells were immunostained with antibodies against FLAG and TOM20. Graphs show the fluorescence intensity profiles of EGFP-LAMP1 and TOM20 along the regions boxed in red. Arrows indicate highly merged LAMP1–CHCHD10^S59L^ region. (**B**) HeLa cells were transfected with siRNAs targeting *NDP52* or *OPTN* or control siRNA. Immunoblotting confirmed RNAi-mediated knockdown of target genes. (**C**) HeLa cells and (**D**) HeLa^YFP-Parkin^ cells were transfected with siRNAs targeting *NDP52* and/or and *OPTN*. Mitochondrial respiration was measured by Seahorse XF Cell Mito Stress tests at 24 hours after transfection.

**Figure S7**

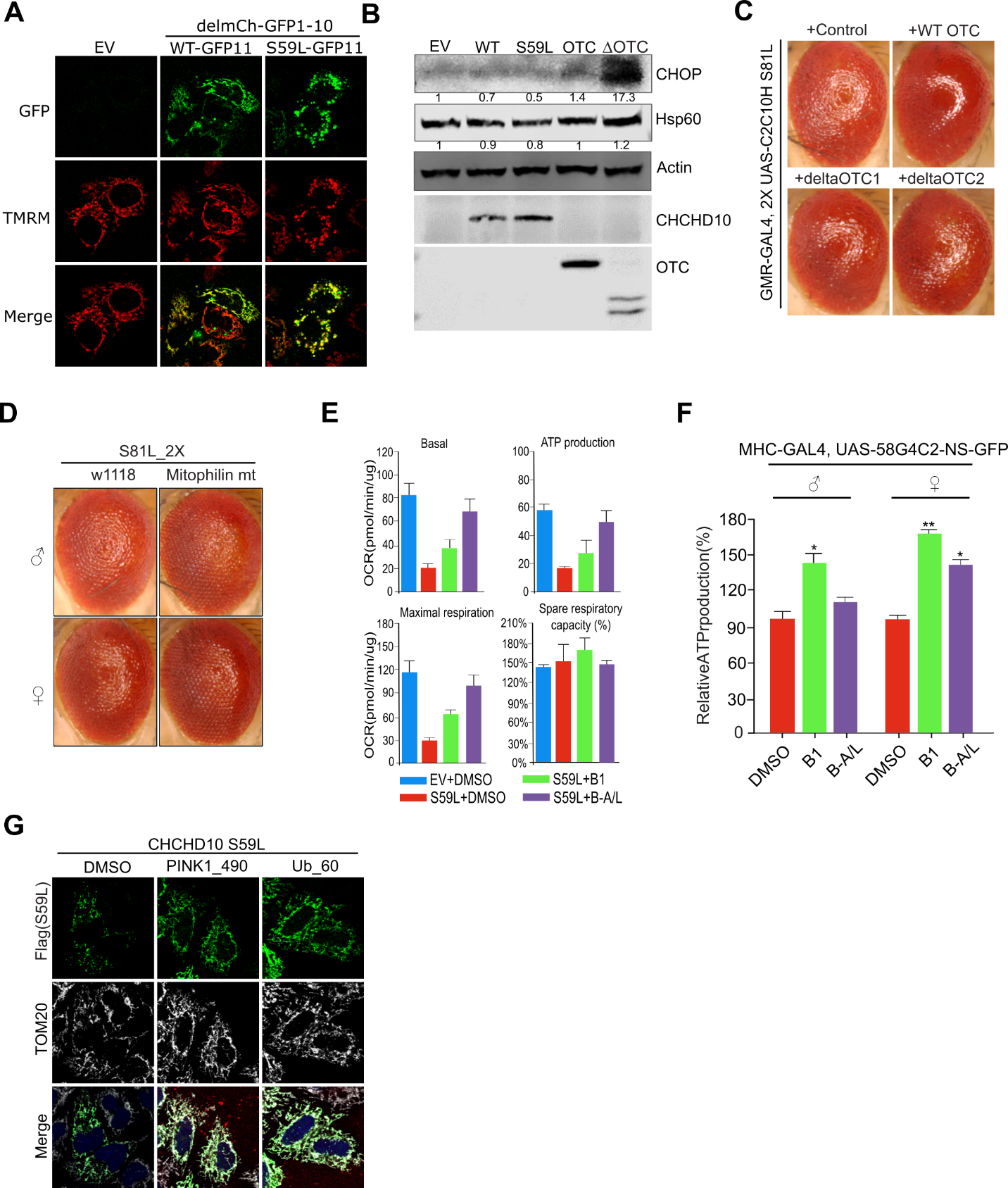

**Fig. S7. Effect of *CHCHD10^S59L^* on mitochondrial membrane potential and the UPR.** (**A**) HeLa cells were transfected with GFP11-tagged *CHCHD10^WT^* or *CHCHD10^S59L^* and mitochondrial-localized GFP1–10. After 24 hours, cells were incubated with TMRM (100 nM). (**B**) HeLa cells transfected with *CHCHD10^WT^*, *CHCHD10^S59L^*, or *OTC* were analyzed by immunoblotting for CHOP and HSP60 proteins. (**C**) Expression of *OTC* or *ΔOTC* did not affect the rough eye phenotype induced by *C2C10H^S81L^*. (**D**) Overexpression of mutant *mitofilin* partially rescued *C2C10H^S81L^*-induced rough eye phenotypes. (**E**) HeLa cells were transfected with FLAG-tagged *CHCHD10^S59L^* and treated with the MFN2 agonists B1 (50 nM) or B-A/L (5 nM) for 24 hours. Mitochondrial respiration was measured with the Seahorse XF Cell Mito Stress test kit. (**F**) ATP levels in thoraxes from 10-day-old flies expressing *C9orf72* with expanded GGGGCC repeats fed with MFN2 agonists (each 10 μM). DMSO (0.1%) was used as a vehicle control. (**G**) HeLa cells were pre-transfected for 24 hours with FLAG-tagged *CHCHD10^S59L^* and then treated with the peptide inhibitors PINK1_490 (0.15 µg/ml) and Ub_60 (0.15 µg/ml). DMSO was used as a vehicle control. HeLa cells were visualized with antibodies against FLAG and TOM20.

**Figure S8**

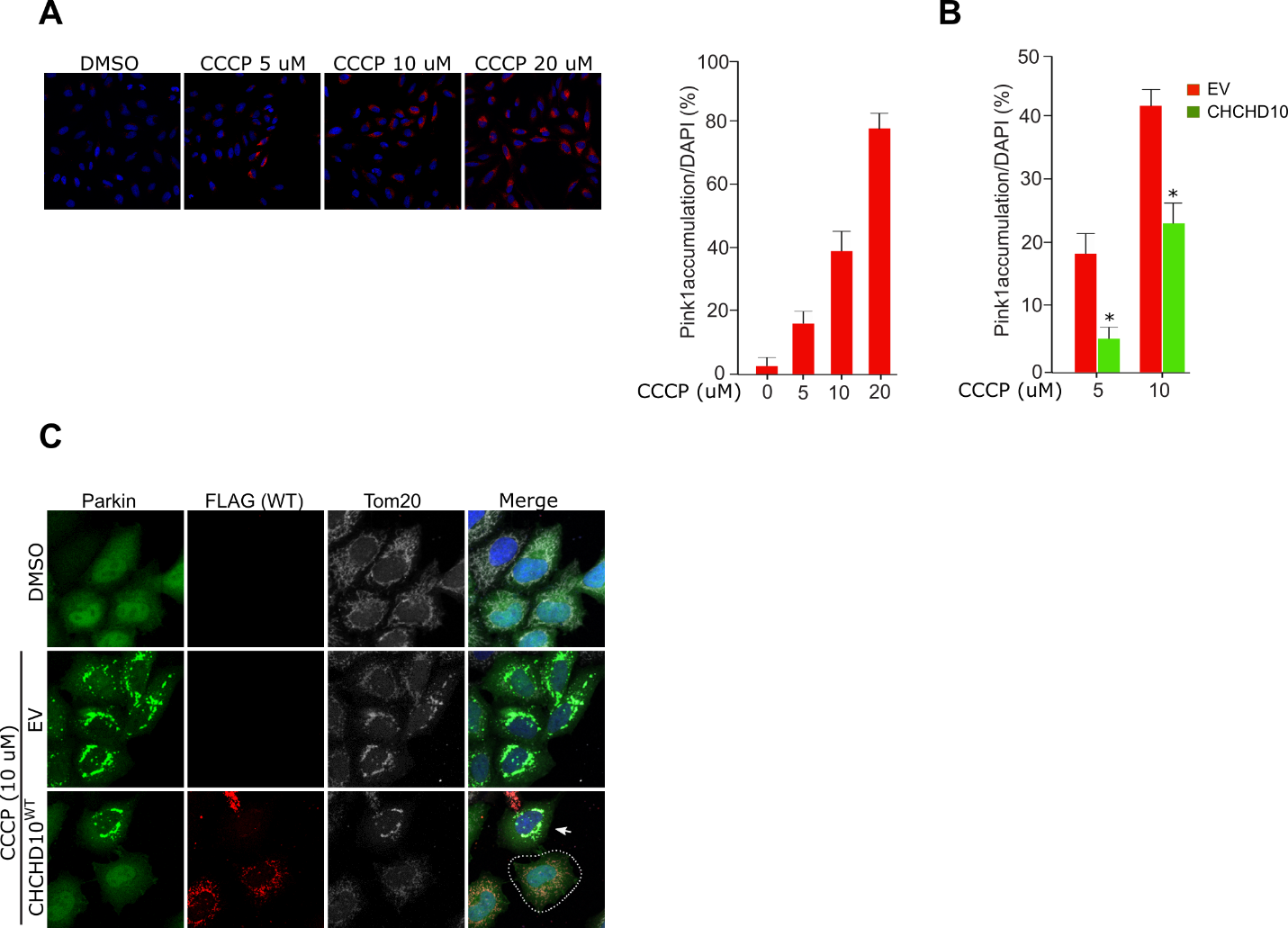

**Fig. S8. Effect of *CHCHD10* on CCCP-induced PINK1/parkin accumulation.** (**A**) HeLa^PINK1-V5-His^ cells treated with multiple concentrations of CCCP for 6 hours were analyzed with a V5 antibody (red) and DAPI (blue). The percentage of PINK1-positive cells was calculated from the number of DAPI^+^ cells. For each sample, at least 300 cells were counted (*n* = 3 biological replicates). (**B**) The percentage of PINK1-positive cells from empty vector (EV)- or *CHCHD10^WT^*-transfected cells were calculated after 5 or 10 μM CCCP treatment for 6 hours. Data are mean ± SD (one-way ANOVA, **P* < 0.05; *n* = 3 independent experiments). (**C**) HeLa^YFP-Parkin^ cells transfected with EV or FLAG-tagged *CHCHD10^WT^* were treated with CCCP (10 μM) for 6 hours. Cells were analyzed with anti-FLAG (red) antibody, YFP (green), and DAPI (blue, nucleus) to visualize CHCHD10 and parkin proteins. Arrow indicates parkin accumulated in nontransfected cell neighboring a *CHCHD10*-transfected cell (white dashed line)

**Table S1. *Drosophila* lines used in study.**

| **Fly Gene** | **Human Gene** | **Stock No.**  **(RNAi or LOF)** | **Stock No. (UAS or Duplicate)** | **Note** |
| --- | --- | --- | --- | --- |
| *Pink1* | *PINK1* | V21860, V109614, B55886, B51650 | *UAS*-dPINK1 (*42*) | RNAi: suppressor,  *UAS*: enhancer |
| *Park* | *PRKN* | V47637, V104363, B38333 | *UAS*-Parkin (*42*) | RNAi: suppressor,  *UAS*: enhancer |
| *Marf* | *MARF* | B31157, B55189, V40478, V105261 | *UAS*-Marf (*42*), B32267, B30293 | RNAi: enhancer,  *UAS*: suppressor |
| *Drp1* | *DNM1L* | B27682, B51483, V44155, V44156 | B51647 | RNAi: enhancer,  *UAS*: enhancer (lethal) |
| *ND-42* | *NDUFA10* |  | B58467 | *UAS*: No effect |
| *sicily* | *NDUFAF6* |  | B58465, B67141 | *UAS*: No effect |
| *Mitofilin* | *IMMT* | V47615, V47616, V106757 | *UAS*-Mitofilin (*48*), *UAS*-MitofilinPR (*48*) | RNAi: enhancer,  *UAS*-PR: suppressor |
| *Miro* | *RHOT1* | B27695, B43973 | *UAS*-dMiro (*60*), *UAS*-Myc-dMiro (*60*) | RNAi: enhancer,  *UAS*: enhancer |
| *ari-1* | *ARIH1* | B29416, B1538 | B30756 | RNAi: No effect,  Duplicate: No effect |
| *mul1* | *MUL1* | V109808 |  | RNAi: No effect |
| *Dmel\CG9855* | *MARCH5* | V105711 |  | RNAi: No effect |
| *TER94* | *VCP* | V24354 | *UAS*-Ter94 (*42*) | Both: enhancer |
| *Atg1* | *ULK1* | B26731, B35177, B44034 | B51654, B51655 | RNAi: enhancer,  *UAS*: enhancer (lethal) |
| *ref(2)P* | *SQSTM1* | B33978, B36111 |  | RNAi: No effect |
| *mib1* | *MIB1* | B27320 |  | RNAi: No effect |
| *mib2* | *MIB1* | B57833 |  | RNAi: No effect |
| *CG5059* | *BNIP3* | B42494 |  | RNAi: No effect |
| *key* | *IKBKG* | B35572, B57759 |  | RNAi: No effect |
| *mud* | *NUMA1* | B28074, B35044, B38190 |  | RNAi: No effect |
| *tango11* | *MFF* | B36281 |  | Deletion mutant: No effect |
| *TBPH* | *TARDBP* |  | *UAS*-hTDP-43 M337V (*38*) | *UAS*-Disease-causing mutant: lethal |
| *TBPH* | *TARDBP* |  | Gene replacement: WT, G294A, and M337V | G294A and M337V: enhancer |
